# Criticality in Tumor Evolution and Clinical Outcome

**DOI:** 10.1101/314021

**Authors:** Erez Persi, Yuri I. Wolf, Mark D.M. Leiserson, Eugene V. Koonin, Eytan Ruppin

## Abstract

How mutation and selection determine the fitness landscape of tumors and hence clinical outcome is an open fundamental question in cancer biology, crucial for the assessment of therapeutic strategies and resistance to treatment. Here we explore the mutation-selection phase-diagram of 6721 primary tumors representing 23 cancer types, by quantifying the overall somatic point mutation load (*ML*) and selection (*dN/dS*) in the entire proteome of each tumor. We show that *ML* strongly correlates with patient survival, revealing two opposing regimes around a critical point. In low *ML* cancers, high number of mutations indicates poor prognosis, whereas high *ML* cancers show the opposite trend, due to mutational meltdown. Although the majority of cancers evolve near neutrality, deviations are observed at extreme *MLs*. Cancers with the highest *ML* evolve under purifying selection, whereas those with the lowest *ML* show signatures of positive selection, demonstrating how selection affects cancer fitness. Moreover, different cancers occupy specific positions on the *ML-dN/dS* plane, revealing a diversity of evolutionary trajectories. These results support and expand the theory of tumor evolution and its non-linear effects on survival.

**Significance Statement:** It remains an open fundamental question how mutation and selection co-determine the course of cancer evolution. We construct a selection-mutation phase diagram, using tumor mutation load and selection strength as key variables, and assess their association with clinical outcome. We demonstrate the existence of a biphasic evolutionary regime, whereby beyond a critical *ML*, the fitness of tumors decreases with the number of mutations, while the proteome evolves near neutrality. Deviations from neutrality in extreme *ML* elucidate how positive and purifying selections maintain tumor fitness. These results empirically corroborate the existence of a critical state in cancer evolution predicted by theory, and have fundamental and likely clinical implications.

## Introduction

The paradigm of tumor clonal evolution by acquisition of multiple mutations has been firmly established since the landmark work of Knudson (1), Cairns (2) and Nowell (3). Similarly to microbial populations (4-6), tumors evolve under constant selective pressure, imposed by the microenvironment as well as by therapy, such that surviving tumor cell lineages harbor mutations that confer selective advantage and resistance to treatment. This has been demonstrated both in space, showing intratumor branched evolution across different anatomical sites (7), and in time, showing the existence of a population bottleneck following treatment, and rapid emergence of resistant phenotypes (8). Under this paradigm, the evolutionary trajectories of cancers can be viewed as different realizations of the same evolutionary process, shaped by the specific microenvironment, the genomic makeup of each tissue and individual, and the unique history of mutations in each clone (3, 9).

Notwithstanding the importance of epigenetics, tumor evolution is marked by a wide range of genomic aberrations and instabilities. These genomic changes occur at every length scale and accumulate in a highly non-linear manner, as exemplified by local elevated mutation rates (Kataegis) (10), complex short insertions and deletions (11), hypermutation and microsatellite instability (12), punctuated equilibrium and chromosomal rearrangements (Chromoplexy) (13), and biased distribution of mutations across different genomic regions (14). Eventually, these somatic aberrations provide for the ability of cancers to proliferate, invade and metastasize (15) by affecting a plethora of cellular functions (16).

Although recent advances in cancer genomics have greatly improved our understanding of how somatic genomic aberrations are linked to tumor progression and patient survival (17-20), the fundamental question how mutation and selection jointly determine the clinical outcome remains open (21-23). The population-genetic theory of tumor evolution predicts that there exists a critical mutation-selection state that corresponds to a transition between evolutionary regimes (24-25). Below the critical state, mutations that increase tumor fitness, known as cancer drivers (26-28), are the main factors of tumor evolution, whereas above the critical state, accumulation of (moderately) deleterious passenger mutations outcompete the drivers, eventually leading to tumor regression through mutational meltdown (25), a process known in population genetics as *Muller’s Ratchet* (29). However, the rarity of spontaneous tumor regression, coupled with strong evidence of increased cancer risk at high mutational loads in hypermutator genotypes (30), contest the existence and relevance of such criticality in clinical outcome.

Furthermore, recent studies indicate that the bulk of cancers and most genes in tumors evolve neutrally (31-33). Conversely, somatic evolution of some normal tissues appears similar to that detected in certain cancers (34), in particular, showing comparable signatures of positive selection (35). Together, these findings prompt the fundamental question how different mutation-selection regimes of tumor evolution determine cancer fitness and ultimately patient survival. Here we address this problem by exploring the dependence of tumor fitness and clinical outcome on mutation load and selection, and demonstrate the existence of criticality in tumor evolution.

## Results

### Population Genetics Approach for Assessing Tumor Evolution and Fitness

To study the inter-relationship between mutation, selection and clinical outcome on a large scale, we quantified the evolutionary state of 6721 primary tumors that represent 23 different cancer types from The Cancer Genome Atlas (TCGA) database (**Methods** and **Figure S1**). The time of tumor initiation and the non-linearity in the accumulation of mutations during its evolution to a primary state are unknown. Further, although the number of cancer-stem cells that confer tumorigenic renewal potential is believed to be small, their actual prevalence and impact on the fitness of tumors remains incompletely understood (36-37). Thus, from the available data that typically present a single snapshot in time of primary tumor states, the effective population size (*Ne*) cannot be reliably determined. Therefore, we define the evolutionary status of each tumor by the overall mutation load (*ML*), i.e. the sum of non-silent (*N*) local somatic genomic alterations including point mutations, small deletions and insertions, and the strength of selection (*dN/dS*), i.e. the ratio of non-synonymous to synonymous nucleotide substitution rates, acting on the entire protein-coding exome (hereafter, proteome) (**Methods** and **Figure S2**).

Respectively, *dN/dS* and *ML* can at least conceptually serve as proxies for the effective population size (*Ne*) and the mutation rate (*μ*), the key variables that are conventionally used in population genetics (21), which determine the evolutionary fates of all organisms (38). This is the case because *dN/dS* and *Ne* are inversely related (39), so that high *Ne* implies dominance of purifying selection, a common evolutionary regime in prokaryotes and unicellular eukaryotes, whereas low *Ne* implies the dominance of neutral evolution by genetic drift, a typical scenario in at least some groups of multicellular eukaryotes (40-41). The case of *ML*, an important clinical measure, is somewhat more complicated. It represents the integration of all non-silent somatic point mutations across the proteome over an unknown but defined time interval. Because some mutations could have accumulated prior to tumor initiation (42), this interval can be defined as the time from the birth of the cell that eventually transformed into a neoplastic cell to the primary tumor state. Thus, *ML* represents the product of the mutation rate and an effective evolutionary time; nonetheless, it can be translated into mutation rate under simplifying assumptions, as we discuss below.

Assuming that the survival of patients is inversely proportional to the fitness of tumors, we explored how *ML* and *dN/dS* correlate with survival, using the semi-parametrized Cox regression analysis and the empirical Kaplan-Meier (KM) log-rank test as two complementary approaches, to increase the significance of the analysis (**Methods**). These tests were applied to both clinical overall survival (OS) and disease free survival (DFS) times.

### Criticality in Clinical Outcome as Function of Mutation Load

First, we explored how *ML* correlates with clinical outcome. To estimate *ML*, we considered all non-silent (*N*) somatic mutations in each patient, including missense (82.3%), in and out of frame insertions and deletions (8.6%), nonsense (5.8%) and splice-site/region (3.2%) variants (**Figure S1**). The distribution of *ML* across the 23 cancer types is in full accord with the well-known ordering of cancers (27-28) in which Thymoma and Acute Myeloid Leukemia (AML) have the lowest *ML*, whereas Lung and Melanoma exhibit the highest *ML* (**Figure 1A,** *top*).

**Figure 1:**
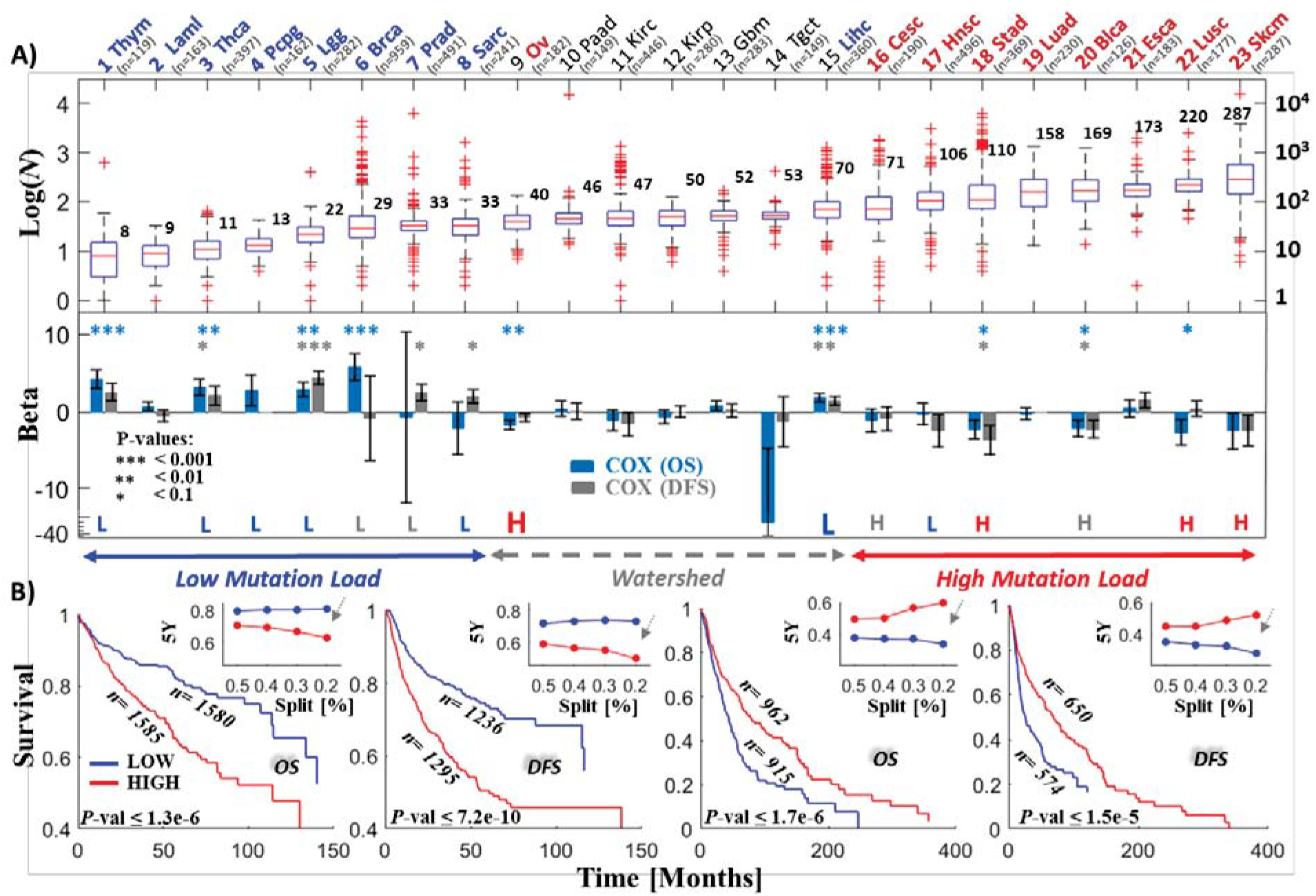
Mutational load criticality in clinical outcome across cancer types. **A)** Log distributions of the number of non-silent (*N*) mutations per proteome/sample in each cancer type; the corresponding results of Cox regression analysis are shown underneath the plot. The KM results (**Figure S3**) are superimposed; the cases where, low (L, blue) or high (H, red) number of mutations was significantly associated with better survival (P≤0.1) are highlighted. Statistical significance is indicated for two additional thresholds, P < 0.01 and P < 0.001. Grey letters (L or H) indicate an observed but not significant (*P>0.1*) correlation. **B)** The KM overall survival (OS) and disease free survival (DFS) rates for sets of cancer types with low mutation load (Left: #1-8 including #15) and high mutation load (Right: #16-23 including #9). For each set, samples with low numbers of mutations (blue; lower 20%) were compared to those with high numbers of mutations (red, upper 20%). Insets depict the 5-years survival rate at different cutoffs (upper/lower percentiles in the range 50%-20%). Arrows indicate the case of ±20% shown in the larger panels. Complementing Cox regression models stratified by cancer types in each group are summarized in **Table 1**. Oncotree codes: **1) Thym**, Thymoma, **2) Laml**, Acute Myeloid Leukemia, **3) Thca**, Thyroid Carcinoma, **4) Pcpg**, Pheochromocytoma and Paraganglioma, **5) Lgg**, Brain Lower Grade Glioma, **6) Brca**, Breast Invasive Carcinoma, **7) Prad**, Prostate Adenocarcinoma, **8) Sarc**, Sarcoma, **9) Ov**, Ovarian Serous Cystadenocarcinoma, **10) Paad**, Pancreatic Adenocarcinoma, **11) Kirc**, Kidney Renal Clear Cell Carcinoma, **12) Kirp**, Kidney Renal Papillary Cell Carcinoma, **13) Gbm**, Glioblastoma, **14) Tgct**, Testicular Germ Cell Cancer, **15) Lihc**, Liver Hepatocellular Carcinoma, **16) Cesc**, Cervical Squamous Cell Carcinoma and Endocervical Adenocarcinoma, **17) Hnsc**, Head and Neck Squamous Cell Carcinoma, **18) Stad**, Stomach Adenocarcinoma, **19) Luad**, Lung Adenocarcinoma, **20) Blca**, Bladder Urothelial Carcinoma, **21) Esca**, Esophageal Carcinoma, **22) Lusc**, Lung Squamous Cell Carcinoma, **23) Skcm**, Skin Cutaneous Melanoma.

We performed a univariate Cox analysis for each cancer type separately. To ensure that the hazard ratios (*HR*) associated with the different *ML* variables are comparable across cancer types, the values of the *ML* within each cancer type were normalized to 0-1 (**Methods**). The Cox analysis of both the OS and DFS of each cancer type reveals two opposing trends of clinical outcome (**Figure 1A,** *bottom*). Among the low *ML* cancers (first 8; median *ML*<40), those that have accumulated higher numbers of *N* mutations, on average, have poorer prognosis than those with lower numbers of *N* mutations (*β*>0, where *β* is the coefficient of the Cox analysis such that *HR*=*e^β^*; see **Methods** for details). However, the relationship between *ML* and survival reverses in high *ML* cancers (last 8; median *ML*>70), where a higher number of *N* mutations corresponds to a better prognosis (*β*<0). Cancers with medium *ML (#9 to #15*) do not show a significant association with survival (*β* ~ 0) except for Ovarian (*#9*, median *ML=40*) and Liver (*#15*, median *ML*=70) at the two sides of the mutation “watershed”, where the pattern of *ML* distributions flattens (*ML* medians ~50). The complementary KM analysis, where we compared the prognosis for patients with low and high *ML* values within each cancer, also captures this transition in the clinical outcome (**Figure 1A** and **Figure S3**). Notably, ovarian cancer behaves as a typical high *ML* cancer type, whereas liver cancer behaves as a low *ML* cancer type, indicating that the mutation watershed represents a critical point in the *ML*-survival dependency. Viewing the flat mutation watershed as a point in *ML*, it is conceivable that cancers in its vicinity can swap positions, such that liver exhibits characteristics of a low *ML* cancer type, whereas ovarian cancer exhibits characteristics of a high *ML* cancer type.

**Figure 1A** depicts a striking overall correlation between the behavior of *β* and *ML* across cancer types. Nonetheless, because the Cox and KM analyses of some individual cancers are not statistically significant, presumably due to the small number of patients, we further tested the existence of opposite regimes, by increasing the statistical power of the analysis (**Figure 1B** and **Table 1**). To this end, we compared between two groups of cancers below and above the watershed. We performed two comparisons of these groups, either including or excluding cancers at the edges of the watershed: **1)** comparing the low *ML* cancers (#1-8) including liver (#15) (**L1**) with the high *ML* cancers (*#16-23*) including Ovarian (*#9*) (**H1**), and **2)** comparing the low *ML* cancers (#1-8) (**L2**) with the high *ML* cancers (*#16-23*) (**H2**), excluding cancer types at the edges of the watershed (i.e., liver and ovarian). The results of the KM tests for the first comparison clearly demonstrate the existence and significance of the transition in clinical outcome (**Figure 1B**). Further, to account for differences between cancer types, we performed complementary Cox regression analyses, in which the data were stratified by the cancer types in each group **(Methods**). The results of this analysis further substantiate the significance and existence of opposing regimes in low versus high *ML* cancers, and demonstrate that the results are robust to the inclusion or exclusion of a particular cancer type in the analysis of either group (**Table 1**).

**Table 1:** Stratified Cox regression analysis of *ML*, overall *CNA, DNA Burden* and *dN/dS* in different cancer groups. For each tested variable, the estimated scaling coefficient *β* (i.e., *HR = e^β^*), its standard error (SE) and the corresponding *P*-value of the stratified Cox regression model are shown for overall survival (OS) and disease free survival (DFS). Significant trends are indicated by bold type and color (*β*>0 blue; *β*<0 red). Cancer groups: Set#1 (L1, H1) corresponds to low *ML* (#1-8) and high *ML* (#16-23) cancer types, including the edges of the mutation watershed (i.e., #15, liver, is included in L1 set, and #9, ovarian, is included in H1 set). Set#2 (L2, H2) excludes these tips. Oncotree codes are as in **Figure 1**. In each test/group, variables are normalized to 0-1, and are stratified by the cancer type (**Methods**).

| Variables (Set) | OS |  | DFS |  |
| --- | --- | --- | --- | --- |
|  | Beta (SE) | P-value | Beta (SE) | P-value |
| *ML* (All) | -1.63 (1.16) | 0.1621 | -1.14 (1.00) | 0.2575 |
| *ML* (L1) | **4.41** (1.02) | **1.62e-05** | **3.12** (0.76) | **3.8e-05** |
| *ML* (H1) | **-4.93** (1.75) | **0.0048** | **-4.24** (1.73) | **0.01452** |
| *ML* (L2) | **3.48** (1.46) | **0.017** | **2.81** (0.89) | **0.0015** |
| *ML* (H2) | **-4.79** (1.73) | **0.0057** | **-4.18** (1.73) | **0.0155** |
| *CNA* (All) | **0.9** (0.19) | **2e-6** | **1** (0.2) | **7.3e-7** |
| *CNA* (L1) | **1.87** (0.3) | **2e-10** | **1.42** (0.28) | **2.7e-7** |
| *CNA* (H1) | 0.31 (0.26) | 0.24 | 0.51 (0.3) | 0.09 |
| *CNA* (L2) | **1.98** (0.31) | **2e-9** | **1.35** (0.32) | **1.8e-5** |
| *CNA* (H2) | 0.27 (0.22) | 0.22 | 0.21 (0.25) | 0.39 |
| *Burden* (All) | **0.48** (0.1) | **2.5e-6** | **0.37** (0.11) | **6.8e-4** |
| *Burden* (L1) | **1.15** (0.19) | **3.3e-9** | **0.84** (0.18) | **1.7e-6** |
| *Burden* (H1) | 0.15 (0.14) | 0.3 | -0.05(0.16) | 0.76 |
| *Burden* (L2) | **1.17** (0.22) | **1.9e-7** | **0.86** (0.21) | **2.9e-5** |
| *Burden* (H2) | 0.14 (0.15) | 0.35 | -0.07(0.17) | 0.71 |
| *dN/dS* (All) | -0.5 (0.4) | 0.21 | -0.12 (0.39) | 0.76 |
| *dN/dS* (L1) | -0.42 (0.52) | 0.43 | -0.38 (0.51) | 0.46 |
| *dN/dS* (H1) | 0.07 (0.52) | 0.89 | 0.42 (0.54) | 0.43 |
| *dN/dS* (L2) | -0.62 (0.55) | 0.26 | -0.54 (0.53) | 0.31 |
| *dN/dS* (H2) | 0.48 (0.58) | 0.41 | 1.08 (0.68) | 0.11 |

### Robustness and Validation of Criticality in Clinical Outcome

To test how robust is the distinction between the opposite cancer evolution regimes with respect to *ML*, we estimated *ML* using different sets of genes, including known cancer-genes and random sets (**Methods**). The emergence of opposite evolutionary regimes around the watershed was highly robust to the choice of the set of genes compared (**Figure S4**). This robustness stems from the high correlation between *ML* values estimated for different sets of genes, which results in similar associations of the *ML* of each set of genes with patients’ survival. Thus, the existence of criticality does not seem to depend on a particular set of mutations or genes, but is rather a consequence of the overall accumulation of diverse mutations in the proteome.

Given that the overall *ML* represents summation over different types of mutational events, it appears likely that other somatic aberrations could provide a comparable signal predictive of survival. Thus, we tested how copy-number alterations (*CNA*) predict survival. We used two standard estimators (linear and gistic) to evaluate the overall *CNA* as well as the overall level of deletions and amplifications in each proteome (**Methods**). We found that *CNA* and *ML* are moderately correlated (Spearman *ρ*=0.44) **(Figure S5)**. However, Cox analysis applied to each cancer type showed that, although at low *ML*, high *CNA* corresponds to poor prognosis (*β*>0), it does not predict the transition in clinical outcome around the mutation watershed **(Figure S5)**. Thus, the Muller’s Ratchet effect at high *ML*, most likely, is caused primarily by point mutations and other small scale mutational events. These observations were confirmed with a stratified Cox analysis comparing low with high *ML* cancers (**Table 1**). Further, we tested the association of the commonly used variable, DNA burden, defined by the fraction of genes affected by *CNA*, finding that it displays similar behavior to the overall *CNA* (**Table 1**). The contrast between the substantial effect of *CNA* in low *ML* cancers and the lack of such effect in high *ML* cancers (**Table 1** and **Figure S5**) suggests nonlinearity, whereby the positive effect of increased *CNA* on tumor fitness is diminished as *ML* increases, consistent with previous findings indicating the association of intermediate copy-number DNA burden values with better prognosis (20).

Testing for the effects of possible confounding factors, including age, stage and grade, by building stratified multivariate Cox regression models (**Methods**), established that *ML* is the only factor responsible for the transition in clinical outcome (**Table S1**). Advanced age and stage, and to a lesser extent grade, were significantly associated with poorer clinical outcome (*β*>0), both in low and high *ML* cancers. However, the transition between the low *ML* cancers (*β*>0) and high ML cancers (*β*<0) was observed only for *ML* (**Table S1**), in agreement with the results shown in **Table 1**.

Lastly, we validated the existence of the transition in clinical outcome by analyzing an independent recent cohort of ~10,000 patients (43) (see **Methods** and **Figure S6**). Although in this data set, only ~400 genes were sequenced, which limits the attainable statistical significance, compared to the TCGA pan-cancer data set, we observed that for low *ML* cancers, the prognostic factor *β* was always positive, whereas in most of the high *ML* cancer types, *β* was negative (**Figure S6**). Thus, the results of this analysis on an extended data set largely recapitulate the transition in clinical outcome as function of *ML.*

### Dominance of Neutral Evolution in the Pan-Cancer Data Set

We next estimated the selection (*dN/dS*) acting on the entire tumor proteome in each patient (**Methods**). Because of the highly variable rates of mutations across a tumor genome and the small overall number of mutations, a conventional direct estimation of selection at the gene level is impossible, unless integration of mutations across patients is permitted **(Figure S2).** Therefore, to explore the potential link between the selection at the patient level (rather than the gene level) to the survival of the respective patient, we estimated the selection that affects the entire proteome in each patient (**Methods** and **Figure S2**). Specifically, we calculated the ratio between the number of non-synonymous mutations per non-synonymous site (*pN*) and the number of synonymous mutations per synonymous site (*pS*) across all genes, considering the proteome (or a large group of genes) as a single sequence. The ratio *pN/pS* was used as a proxy for selection (*dN/dS*). In cancer, *pN/pS* is a valid approximation of *dN/dS*, assuming that a site is not mutated more than once during tumor evolution, such that correction for multiple mutations that effectively transforms *pN/pS* into *dN/dS*, is unnecessary (**Methods**).

Estimation of the number of mutations in the entire proteome of each patient shows that, in accord with many previous observations on evolving organisms (44), the numbers of non-silent (*N*) and silent (*S*) mutations are highly correlated and display a linear relationship, albeit with different ratios across cancer types, suggesting some diversity of evolutionary regimes (**Figure 2A**). To ensure that our estimate yielded a stable measure of selection, characteristic of the diversity among cancer types, we examined the dependency of *dN/dS* on the number of genes used for the estimation. The median *dN/dS* value in each cancer type reached a plateau rapidly as more genes were included, and the variance across patients in each cancer type was low (**Figure 2B**). Thus, the median *dN/dS* across an entire proteome appears to be an adequate measure for a pan-cancer comparative analysis. The distributions of *dN/dS* indicate a (near) neutral evolutionary regime, where for most cancer types, *dN/dS* values were distributed around 1 across patients (**Figures 2B** and **2C**). This observation was robust to using only missense point substitutions, instead of all non-silent mutations, for the *dN/dS* estimation (**Figure S7**).

**Figure 2:**
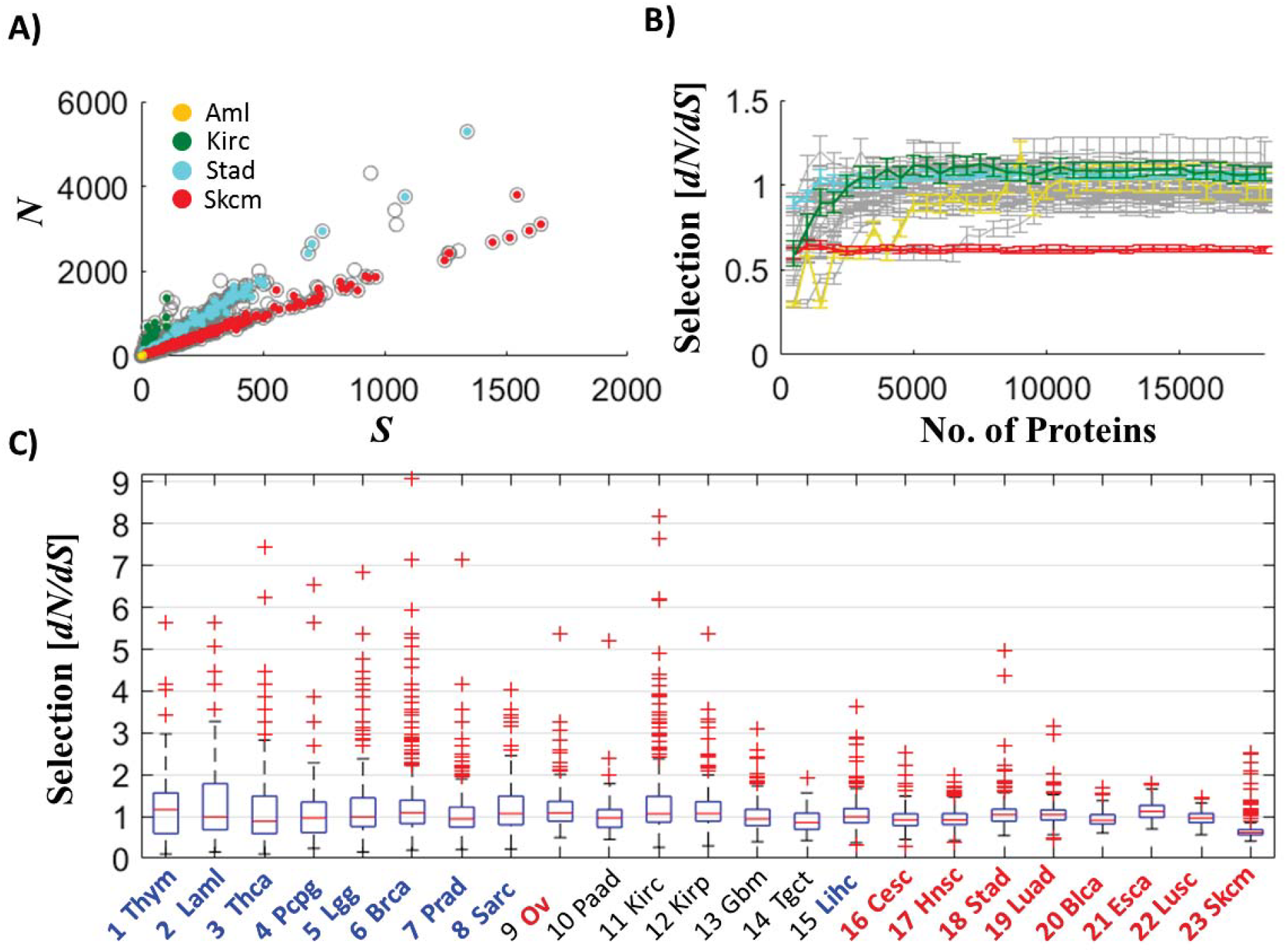
Proteomic selection (*dN/dS*) across cancer types. **A)** The relationship between the numbers of non-silent (*N*) and silent (*S*) mutations per tumor proteome, where representative cancer types that span the different mutational load regimes are color-coded. **B)** Stability of the proteomic measure of selection for comparative analysis between cancer types. The median of protein-level selection (*dN/dS*) across patients is shown as function of the number of proteins considered for the evaluation of *dN/dS*, in each cancer type (grey). Selected cancer types are highlighted as in (A).**C)** Distributions of *dN/dS* in the tumor proteome across patients, for different cancer types.

This result is consistent with those of three recent studies, each using a different approach to estimate selection in tumors (and genes), but all coming to similar conclusions on the prevalence of neutral evolution in the pan-cancer data: (i) an integrative approach which fits the distribution of subclonal mutations in each patient to a *1/f* power law model, by accurate calling of the allele frequencies (*f*) (31), an integrative approach which infers the selection acting on genes, by a applying a Bayesian framework to the overall distribution of mutations (32), and (iii) inference of the exact substitutions rates in different mutational contexts, using a model with 192 parameters (33). Although some differences exist among the methods and conclusions of these studies (see **Methods**), all of them show that the majority of tumors (and genes) evolve close to neutrality. The convergence of all these studies on the predominantly neutral regime of tumor evolution additionally indicates that, at least at the entire proteome level, measures of selection capturing neutral evolution are insensitive to the exact characteristics of mutations (e.g., clonal vs. subclonal) or the distinct (non-linear) dynamics by which different mutations accumulate in the proteome (e.g., variable substitution rate and allele frequency).

### Deviations from Neutrality at Extreme Mutation Loads

Notwithstanding the prevalence of neutral evolution (*dN/dS*~1), **Figure 2** also reveals deviations from neutrality at extreme mutation loads. In Thymoma, the cancer type with the lowest *ML*, the median of *dN/dS* is greater than 1, and more generally, heavier tails of *dN/dS*>1 are observed in low *ML* but not in high *ML* cancers, indicative of positive selection at low *ML*. In contrast, in Melanoma, the cancer type with the highest *ML, dN/dS* was distributed completely below 1 (except for a few patients), which is indicative of purifying selection acting on the tumor proteome. These observations were robust to using only missense point substitutions (**Figure S7**).

To elucidate how these deviations from neutrality emerge across the proteome and to assess their significance, we performed a detailed inspection of the distribution of mutations, across different groups of genes, in Acute Myeloid Leukemia (AML) (**Figure 3A**) and Melanoma (**Figure 3B**), which represent the cancer types with extreme *ML* values. AML was selected as an example of a low *ML* cancer to analyze the heavy tails that are indicative of positive selection although on average it appears to evolve neutrally. The analysis of AML patients (n=163) shows that 64 patients had *dN/dS* ≥ *1*, and 63 had *dN/dS<1* **(Figure 3A)**, leading to the observed median of *dN/dS*=1. The remaining 36 patients harbored many *N* mutations, but not a single *S* mutation (i.e., *dN/dS*=*Inf*, which is discarded from analysis); hence, the heavy tail in AML patients (cf. **Figure 2C**) is underestimated. The signature of positive selection (*dN/dS*>1), manifested by heavy tails of the *dN/dS* distributions, was detected in AML samples that harbored numerous mutations, and therefore could not be an artifact caused by the small number of mutations in low *ML* cancers. Furthermore, *dN/dS*<1 in AML patients was a consequence of the large number of *S* mutations (and not of increased statistical power). In contrast, in the case of Melanoma, *dN/dS* values were below unity in the vast majority of samples, and sharply dropped with the increasing number of mutations in the proteome, in a clear sign of purifying selection correlated with the *ML* (**Figure 3B**).

**Figure 3:**
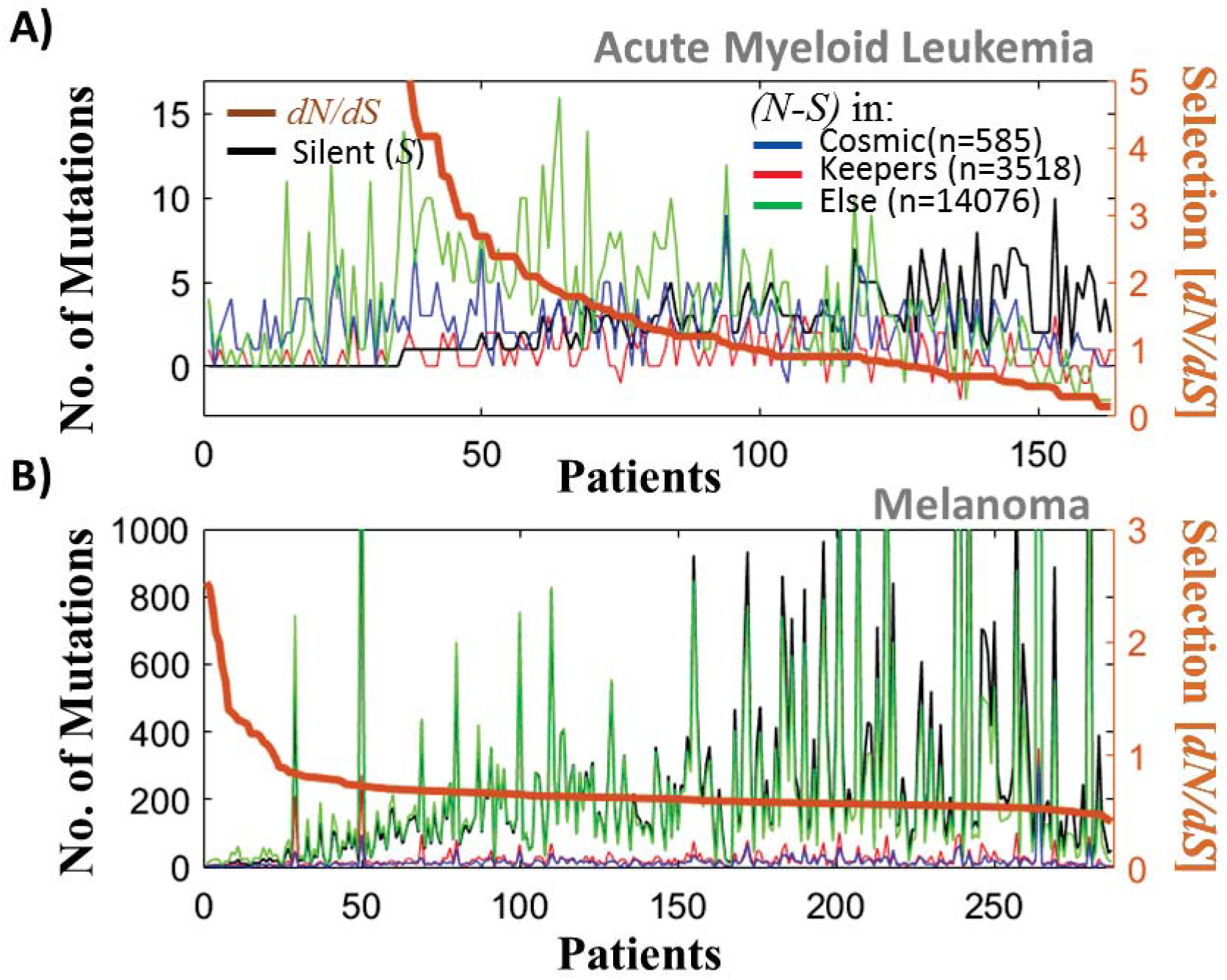
Distribution of mutations in different groups of genes, in cancer with extreme *ML* values. **A)** Acute Myeloid Leukemia patients (n=163). The number of non-silent (*N*) minus the number of silent (*S*) mutations (left y-axis) indicates the excess of *N* mutations in each group of genes separately (color). The number of *S* mutations in the entire proteome is superimposed (black). Patients are ordered by the *dN/dS* acting on the proteome (right y-axis). **B)** Melanoma patients (n=287). In Aml, for more than half of the patients, *dN/dS*>1, and cancer genes harbor a substantial fraction of the *N* mutations. In Melanoma, *dN/dS* is below unity in the vast majority of patients, and *dN/dS* sharply drops with the number of mutations in the proteome which, coupled with *β*<0, indicates mutation meltdown (*Muller’s Ratchet*).

To assess the evolutionary pressures that affect different classes of genes in tumors, we compared the *dN/dS* distributions for the known cancer genes (26) (n=585) and house-keeping genes (45) (n=3518) (**Methods**). The results of this analysis could not be as significant as those for all genes, due to the relatively small number of genes in each set (especially, the cancer genes). Despite this limitation, *dN/dS* in the cancer genes across all cancer types was significantly higher than in randomly selected genes, which was not the case for the house-keeping genes (**Figure S8**). Thus, the cancer genes appear to be subject to stronger than average positive selection. Nonetheless, the accumulation of many *N* mutations outside of the set of known cancer genes indicates that positive selection can affect diverse genes in tumor, with the implication that many cancer-related genes remain to be discovered. In contrast, in Melanoma, purifying selection (*dN/dS*<1) was found to act on large portions of the proteome **(Figure S8).** This signature of purifying selection reflects the fact that, as the *ML* increases, the number of *S* mutations grows faster than the number of *N* mutations across the proteome (**Figure 3B**). Coupled with the observation of better prognosis (*β*<0) in these Melanoma patients (cf. **Figure 1A**), this expansion of mutations across the proteome appears to be a sign of a looming mutational meltdown.

Proteomic measures of selection can provide information on the evolutionary regimes of different groups of genes but not of individual genes (**Methods**). Nevertheless, the results of our analysis are concordant with previous findings (32), showing that in AML more genes are subject to positive than to purifying selection, whereas in Melanoma, the opposite is the case. Furthermore, in Melanoma, the number of genes under purifying selection was found to be greater than in any other cancer type.

### Clinical Outcome Weakly Depends on Selection

To determine whether any of the selection regimes in tumors affect survival, we tested the potential link between *dN/dS* and prognosis. First, we performed KM analysis in each cancer type, comparing positive vs. purifying selection (**Figure S9**). All of these tests failed to detect a significant predictive signal of differential survival. A complementary Cox regression, comparing between the two groups of cancer types with low and high *ML*, stratifying the data by cancer types in each group, verified the lack of association of purifying or positive selection at the proteome level with clinical outcome (**Table 1**). Nonetheless, KM analysis shows that, in certain cancer types (Gbm, Cesc, Lusc, Skcm), intermediate values of selection around neutrality (*dN/dS*~1) were associated with poorer prognosis than either positive or purifying selection (**Figure S10**). Indeed, neutral evolution was associated with poorer prognosis than either type of selection when the comparison was performed across all cancer types although this connection was less significant for disease-free survival (**Figure 4**).

**Figure 4:**
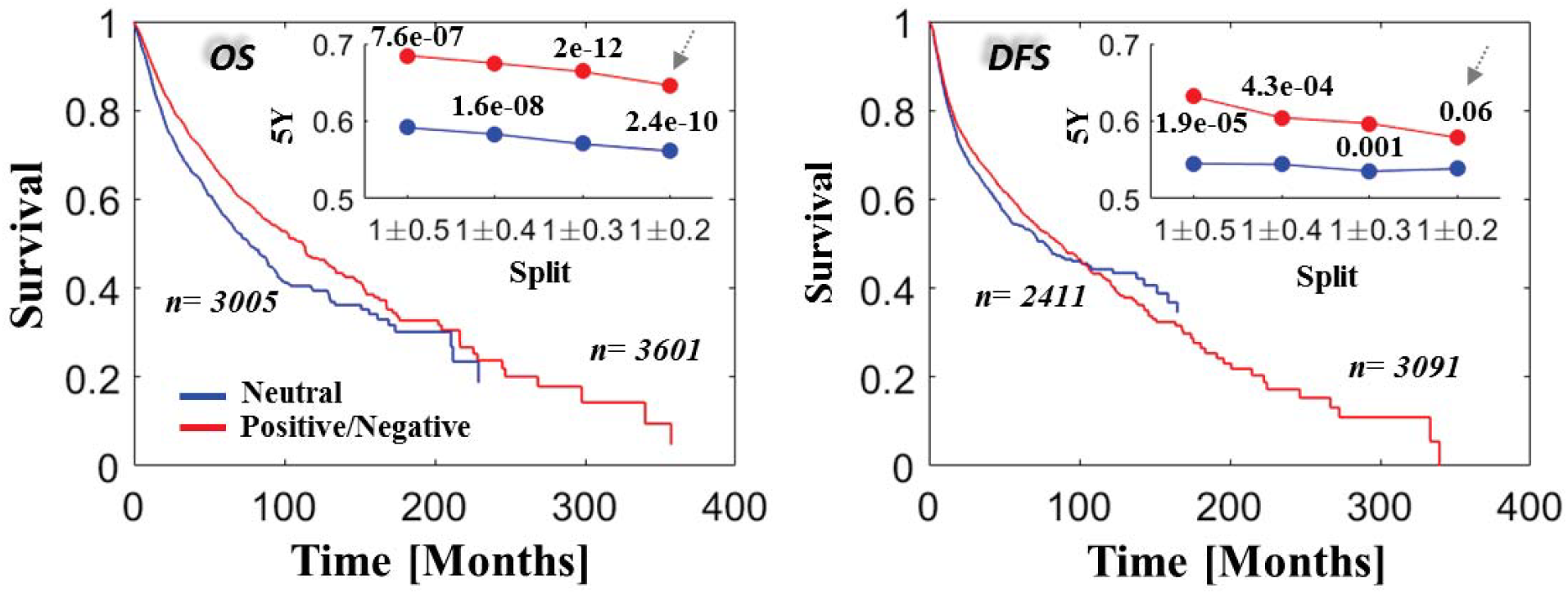
Selection versus survival in the pan-cancer data. KM overall survival (OS) and disease free survival (DFS) rates are compared across all studies for cases of neutral evolution (intermediate values around *dN/dS*=1, blue) and cases of positive and negative selection (red). Insets depict the 5-years survival rates and the corresponding *P*-values of log-rank tests for each cutoff. The survival curves in the larger panels correspond to the case of *dN/dS*=1*±*0.2 as indicated by the arrows in the insets. Complementary Cox regression analysis, stratifying the by cancer types, is provided in **Table 1**.

## Discussion

The results of the present analysis can be best interpreted by projecting *ML* and *dN/dS* onto an empirical mutation-selection phase-diagram, which emphasizes the existence of distinct evolutionary regimes (**Figure 5A**). Under the assumption that cancer fitness and patient survival are inversely related, this diagram shows how *ML* and *dN/dS* jointly determine cancer fitness (**Figure 5B**). In low *ML* cancer types, tumor fitness increases with the number of mutations (*β*>0). In this regime, some tumors appear not to have acquired a sufficient number of driver mutations, and therefore, positive selection (*dN/dS*>1) promotes driver mutations to increase or maintain the tumor fitness (e.g., Acute Myeloid Leukemia). In contrast, at high *ML*, cancer fitness decreases with the number of mutations (*β*<0), due to the accumulation of deleterious passenger mutations. In the cases of extremely high *ML*, this expansion of mutations can lower the fitness of tumors, such that purifying selection (*dN/dS*<1) acts to remove deleterious mutations, thus avoiding tumor collapse by mutational meltdown (*Muller’s ratchet*), as we observed in Melanoma. In Melanoma, *dN/dS* is below unity in samples with large *ML* but turns towards unity in patients with lower *ML* (**Figure 5A**) that on average have a worse prognosis (**Figure S10**).

**Figure 5:**
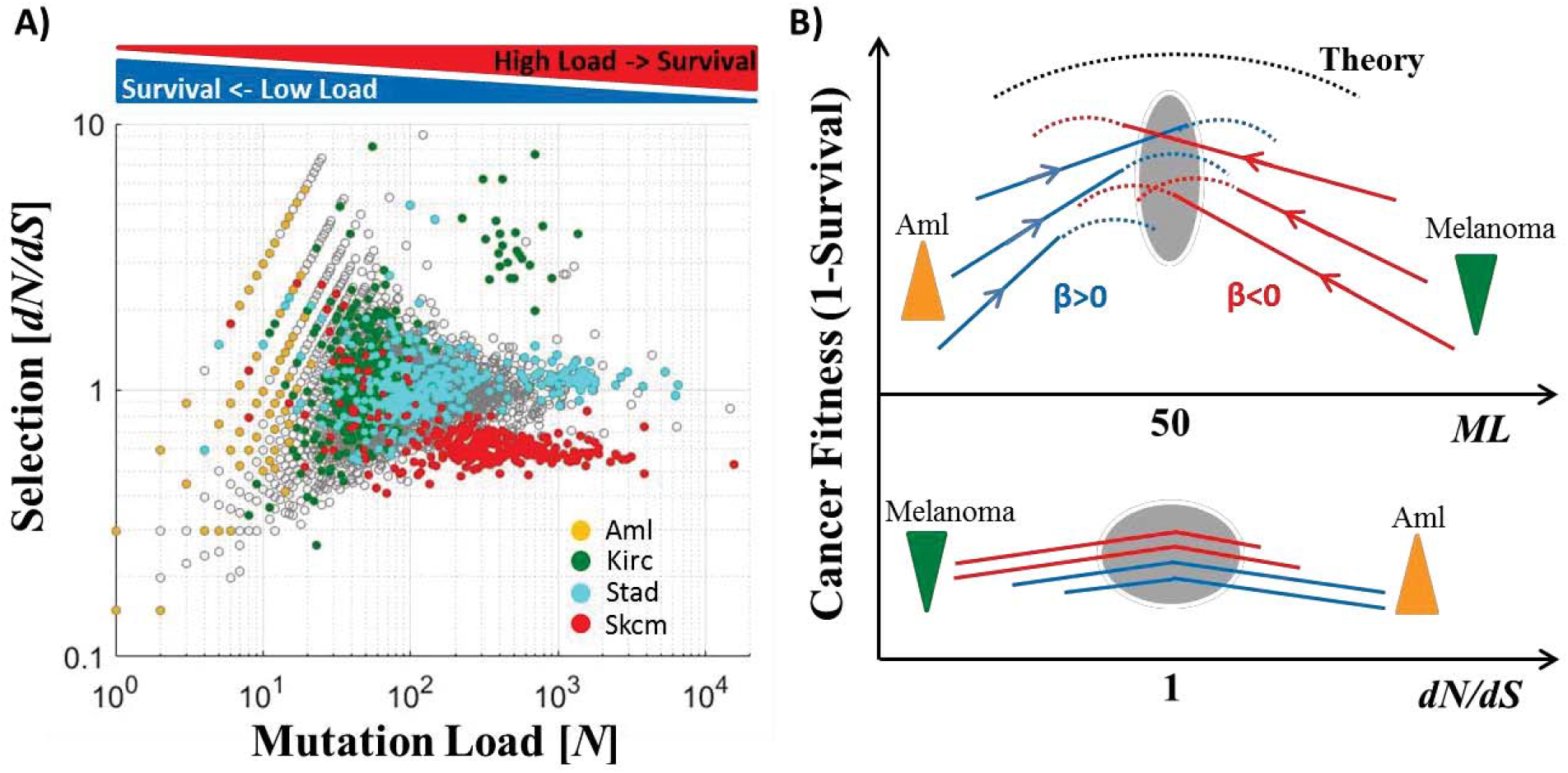
Empirical mutation-selection phase diagram of tumor evolution. **A)** Mutation-selection empirical diagram for all analyzed cancers (grey) and selected cancer types (color-coded) that show distinct evolutionary regimes depending on the mutational load. **B)** A schematic conceptual depiction of the emerging fitness landscape of tumors as function of the mutation load (top) and selection (bottom). Dashed curves are theoretical and solid curves are observed. Down-triangles (green) indicate purifying selection and up-triangles (orange) positive selection. The grey ovals show the critical area.

In contrast to the clear dependency on *ML*, tumor fitness is only weakly correlated with *dN/dS*, such that the majority of cancers evolve near neutrality (**Figure 2**), consistent with previous findings (31-33). This lack of detectable proteomic-level selection signatures is likely due to the fact that tumor fitness mostly depends on a small number of drivers, whereas the bulk of the fixed mutations are neutral or slightly deleterious. Indeed, more detailed analysis (**Figure 3** and **Figure S8**) demonstrated significant differences in selection between groups of genes, in particular, positive selection in cancer genes, with an overall neutral effect on the entire proteome. Only at extremely high *ML*, as in Melanoma, tumor fitness depends on the accumulation of a critical mass of (deleterious) mutations across the entire proteome, so that these tumors evolve under purifying selection. Thus, in summary, under neutrality, a sufficient number of drivers can accumulate whereas the overall deleterious effect of passengers is balanced, explaining the weak association of neutrality with poor prognosis (**Figure 4**). Taken together, our results corroborate the theory of tumor evolution that predicts the existence of a critical mutation-selection state (24-25). Nonetheless, the existence of tumors with high *ML*, some of them with poor prognosis, suggests that other somatic aberrations could increase or maintain tumor fitness, to compensate for the passengers deleterious effect. This seems to be the case of microsatellite instability. In many hypermutation tumors, microsatellite instability is associated with better prognosis, thus apparently reducing tumor fitness (12). Further, high *ML* tumors across different cancer types on average have low microsatellite instability (46). Thus, a compensatory relationship appears to exist between point mutations and microsatellite instability with respect to tumor fitness.

In addition to these general trends, examination of the empirical *dN/dS-ML* plane reveals a diversity of tumor evolution regimes. For example, in kidney renal clear cell carcinoma, we identified a cluster of patients with high *ML* and *dN/dS*>1, suggesting that the specific microenvironment and other factors, such as competition between subclones (21, 47), could be important for understanding the precise relationship between *ML*, *dN/dS* and survival. Hence, coupled with the overall weak association of selection with survival, selection appears to maintain cancer fitness in diverse microenvironmental conditions, genomic contexts and phases of evolution, leading to a diversity of roughly equally successful evolutionary strategies (with respect to *dN/dS*) of extant cancers, while the neutral evolutionary regime dominates overall.

Our analyses indicate that the overall mutational load is a key determinant of patient survival. The *ML* counts all *N* mutations, wherever they occur in the tumor genome (including portions involved in structural variation, such as gene duplication), and whenever they emerge during the lifetime of tumor cells. Given that the survival dependency on *ML* captures the transition in the clinical outcome, the effects of various mutations appear to be context-dependent, and in a given genomic state could lead to either an increase or a decrease in tumor fitness, such that all mutations should be included to assess clinical outcome. Therefore, the total *ML* becomes a key variable for clinical assessment, which is not sensitive to cellularity, ploidy, clonality and other specific features of tumors. The high correlations between *ML* of different classes of genes (**Figure S4**) as well as between *ML* values for different mutation classes (**Figure 2A** and **Figure S7**), with all these values being tissue-specific as in (27-28), suggest that *ML* is a stable measure that reflects effective (tissue-specific) evolutionary time of a tumor (weighted by the respective variable mutation rates). This is consistent with recent observations showing that the tissue-specific cell division rate is a key determinant of cancer risk and the mutational load in diverse tissues, whereby about 2/3 of the mutations accumulate at random due to replication errors (48-49). This is also consistent with the observation that both genetic and epigenetic characteristics of the original cell are key determinates of the mutational spectrum of the respective cancer cell (50).

The criticality observed around the mutation watershed corresponds to the transition in the clinical outcome at *ML* of ~50 *N* mutations per tumor proteome. Under certain simplifying assumptions, this value can be linked to previous results. Data-driven theoretical studies suggest that, for ~60 passengers (*P*=*N*+*S*−*D*; *P*, total number of passenger mutations; *D*, number of drivers among the *N* mutations), there are ~10 drivers (24). Thus, for the critical point as identified here, *N~50, S*~20 and *D*~10. To accumulate 10 drivers, it takes ~5-50 years with a cell division rate of ~4 days (i.e., the number of cell generations *G*=450−4500) (24). Thus, we can estimate that the range of mutation rates (per locus per cell division) associated with *N*~50 is *μ~5×10^−9^−5×10^−10^(μ = N/Ns/G; Ns*, the total number of *N* sites in the proteome). This range of mutation rates closely matches the lower range of rates where a non-monotonic accumulation of passengers vs. drivers starts to be detectable, leading to the effect of Muller’s Ratchet predicted by theory (25). Further, if *D*~10 and each clone in a tumor harbors a small number of drivers (~*2-3*), then the critical number of clones for tumor progression is ~*3-4*, in agreement with recent findings (20).

### Concluding remarks

To summarize, in addition to known genomic markers (18, 20), our results reveal major, global features of cancer genome evolution that affect tumor fitness and accordingly, clinical outcome. In accord with theoretical predictions, we show that the dependency of tumor fitness on the mutational load is non-monotonic, with a critical region where the evolutionary regime changes, empirically corroborating the theory of tumor evolution, as a tag of war between driver and passenger mutations (25). In contrast, the dependency of tumor fitness on proteome-level selection is weak. We conclude that tumor fitness and clinical outcome strongly depend on the total *ML* and that most tumors evolve under a predominantly neutral regime, with relatively small contributions of both purifying and positive selection that become stronger only at extreme *ML* values. These conclusions are compatible with the well accepted view that tumors evolve and progress via random accumulation of a few driver mutations.

By analyzing proteomes of a broad range of cancers, we identify tumors that evolve in different regimes that are characterized by opposite effects of *ML*. Knowledge of the evolutionary status of a given tumor could have implications for therapy that would aim to either increase or decrease the *ML*, depending on the position of the given tumor on the dependency curve. This might be particularly important for immunotherapy, where *ML* plays a critical role (51). Our results further imply that targeted therapy can be effective in low *ML*, where few drivers determine the course of tumor evolution, whereas at high *ML*, alternative strategies, such as immunotherapy, are likely to be more effective, consistent with the well-known success of immunotherapy in melanoma (52-53). The present analysis could also serve as a framework for future research to study how the transition from the primary to the metastatic state and how therapy could change the status of tumors in the *ML-dN/dS-β* hyperplane.

## Materials and Methods

### Datasets

The complete raw data form all TCGA studies (n=23) that included at least 100 patients each were downloaded from cBioPortal (54) (http://www.cbioportal.org/). For analysis, we considered all “3-way complete” samples (i.e., containing somatic point mutations, copy-number alterations and gene expressions data, relative to matched-normal samples; n=6721), and all human protein-coding genes for which we identified both SwissProt and NCBI-Entrez unique accessions (n=18179). This data matrix (samples by genes) as well as patients’ clinical data were also downloaded from Firehorse (https://gdac.broadinstitute.org/) for comparison, verifying that there is little discrepancy between the two databases and that each mutation had at least 10 reads of the tumor variant (standard quality control) and are fully non-redundant (i.e., a variant in a given sample and gene are not counted more than once). Data from cBioPortal were downloaded also via Matlab application program interface (API), which routinely updates annotations of mutations, and were used to remove germline mutations from analysis. Clinical survival data included overall survival (OS) for 98.3% of the patients (n= 6609) and disease free survival (DFS) for 82% of the patients (n=5508). Distribution of patients’ race and age, tumor stage and grade as well as the distribution of variants across different mutational classes are provided in **Figure S1**.

Known cancer genes were downloaded from COSMIC database (26) (http://cancer.sanger.ac.uk/census). House-peeping genes were extracted from a recent survey (45). For validation (**Figure S6**), a recent cohort of ~10000 patients with advanced cancer (MSK-impact-2017), where 43% of the samples originate from metastatic sites and 414 cancer genes were sequenced (43), was downloaded via cBioPortal. Data for all samples and genes, including all the information needed for full reproducibility of the results in this study, are provided in **Supplemental Dataset S1** (Excel file).

### Copy-number alterations (CNA)

To estimate gene copy-number alteration (CNA), we extracted and analyzed both the ‘linear’ and ‘gistic’ measures. Linear measures provide continuous variables which represent the extent of amplification and deletions of each gene. The gistic measure implements additional computation inferencing the zygotic gain/loss using integers (*-2* to *2*). For evaluation of the overall level of CNA (**Table 1, Table S1**), we used summation over the ‘linear’ measure, verifying that it correlated with the summation over the ‘gistic’ values (**Figure S5**). The copy-number DNA burden was also calculated, using the ‘gistic’ measure, as the fraction of altered genes (gain or loss) in the proteome (**Table 1**).

### Selection in Tumor Proteomes

Protein-level selection (*dN/dS*) at the molecular level is measured by comparing two sequences and computing the ratio between the non-synonymous substitution rate (*dN*) and the synonymous substitution rate (*dS*) (55). Generally, this is done in two steps: (i) calculating the number of *N* sites (*nN*) and the number of *S* sites (*nS*) over the length of the compared sequences, and calculating the number *N* mutations per *N* sites (*pN=N/nN*) and the number of *S* mutations per *S* sites (*pS=S/nS*), and (ii) applying methods, such as Jukes-Cantor (56) or Goldman & Yang (57) that transform the counts *pN* and *pS* into the respective rates *dN* and *dS*, by considering the possibility that over time, a single locus mutates several times before fixation, in a context-dependent manner. Over long evolutionary distances, this second step is crucial. During cancer evolution, however, the likelihood for a particular locus to mutate more than once is low (9) and a considerable number of mutations might not be fixed, such that estimates of selections should be based on the integration of mutation counts rather than rates (58). Hence, we chose to approximate *dN/dS* by the ratio *pN/pS*.

Selection can be assigned and computed at different length scales (e.g., locus, domain, gene, etc.). In practice, the pan-cancer mutation data are highly sparse such that a gene in a patient rarely harbors both *N* and *S* mutations (**Figure S2**). Thus, a direct estimation of *dN/dS* at the gene level is not feasible, and integration of mutations, either over patients providing estimates of selection in individual genes, or over genes, providing estimates of selection in individual patients, is necessary. Adequate measures of selection at the gene level have been recently developed, using both a Bayesian framework (32) and a context-dependent inference of substitution rates (33). Here, our goal was to investigate the link between the selection acting on the tumor proteome and the respective patient survival, so data from different patients should not be integrated. Therefore, we compute selection at the patient level, integrating mutations over genes (*g*) within in patient’s tumor proteome and treating them as a single concatenated sequence, such that there are sufficient numbers of *N* and *S* mutations for statistical inference of *dN/dS*:

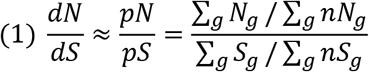

The *dN/dS* values were estimated using in **Equation 1**, for each patient, considering the mutations in the entire proteome (**Figure 2**), or groups of genes, such as known cancer genes or house-keeping genes (**Figure S8**). Practically, to calculate the *dN/dS* ratios, the canonical amino-acid sequences of all human proteins and their respective DNA coding sequences were extracted primarily from Ensembl (59) and from GeneBank for completeness. For each nucleotide sequence, translation into the exact respective canonical protein sequence in SwissProt was verified. The numbers of non-synonymous and synonymous sites (*nN, nS*) in each protein were calculated, considering all alternative nucleotides in each position. The full accord of the selection in entire proteomes (**Figure 2**) with previous studies (31-33), capturing the dominance of neutral evolution, independently validates the choice of **Equation 1** for the large-scale comparative analysis across patients cancer types.

### Survival Analysis

To test the association of variables with survival, we used both Kaplan-Meier (KM) log-rank test (60-61) and Cox proportional hazard regression analysis (62), and applied these approaches to both OS and DFS clinical data. KM is a non-parameterized empirical test that compares the survival curves using long-rank test for censored data. In this analysis, groups of patients are defined and compared by splitting the tested variable. This approach allows flexibility in defining and testing different ranges of the tested parameter, albeit at the risk of losing robustness. Hence, to assess the stability of this test, we used several cutoffs as indicated for each analysis. Cox regression is a semi-parameterized approach that fits the survival clinical data to a hazard function (*h(t*)=−*d[logS(t)]/dt*, where *S(t*) is the survival probability at time *t*) and tests the effect of variables (*X*) under the ‘proportional hazard’ assumption (*h(X,t*)=*h_o_(t)e^Xβ^*; *h_o_* the baseline hazard), namely, that the tested hazard functions are log-linearly scaled by a constant factor beta (*β*), which determines the Hazard ratio (i.e., *HR*=*e^β^*). This assumption, however, does not always hold for real data. Hence, the KM and Cox analyses are complementary.

Using Cox analysis, we normalized each tested variable (e.g., *ML, dN/dS, CNA*) in each test to 0-1, such that the results of different tests can be easily compared, (see also Ref: **20**). Hence, in **Figure 1A**, *ML* is normalized in each cancer type to 0-1, and a univariate Cox analysis is performed in each cancer type separately. Similarly, when several cancer types were grouped (e.g., low or high *ML* in **Table 1**), the aggregated distribution of the *MLs* across patients in each group was normalized to 0-1, and the variables were stratified by the cancer types, to build stratified regression models for each group separately.

To build stratified multivariate regression models (**Table S1**), testing the effects of possible confounding factors such as age, stage and grade, the categorical clinical data, stages I-IV and grades I-IV, were tested each using dummy indicator variables, relative to the reference category stage/grade I, respectively. Subcategories were grouped (e.g., Stages IA-IC were assigned Stage I). Any stage or grade outside the range I-IV (e.g., stage/grade ‘X’) were not included in this analysis, and were not given any value (i.e., Nan). Variables were stratified by cancer types. The constants of each Cox proportional hazard regression model (*β*, its error and the *P*-value) are provided in each figure and table for each test.

### Analysis and Code Availability

All the analyses were performed in Matlab R2016b, using only built-in functions, under license to UMD/UMIACS/CBCB. Matlab files, including the datasets and analysis scripts, which fully reproduce the results as they appear in the manuscript, are available upon request from the authors.

## Acknowledgements

We thank the Koonin group at the NIH for discussions and feedback, and Michael F. Berger for sharing data of the large cohort used for validation.

## Supporting Information

**Figure S1:**
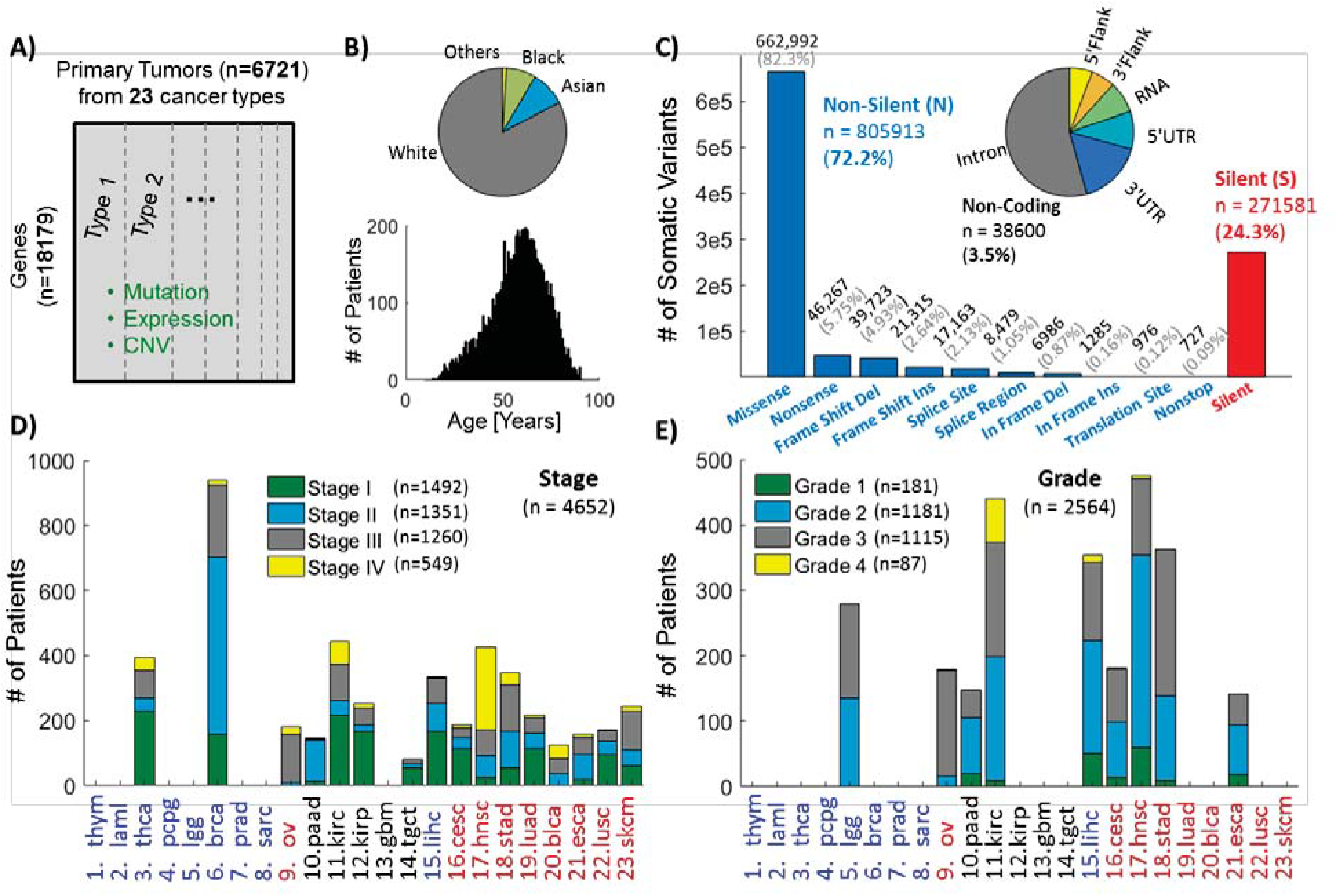
Data Statistics: Primary tumor data downloaded from cBioPortal of all TCGA cancer types containing at least 100 patients each, covering 6721 patients across 23 cancer types. **A)** We analyzed all “3-way complete” samples, for which gene expression, CNV and somatic mutations data exist, and considered all protein coding genes that have both unique NCBI-Entrez and SwissProt IDs (n=18179). **B)** Distribution of race and age across patients. **C)** Distribution of the number of mutations in the proteome associated with each mutation class across patients. **D)** Distribution of stage across cancer types. **E)** Distribution of grade across cancer types.

**Figure S2:**
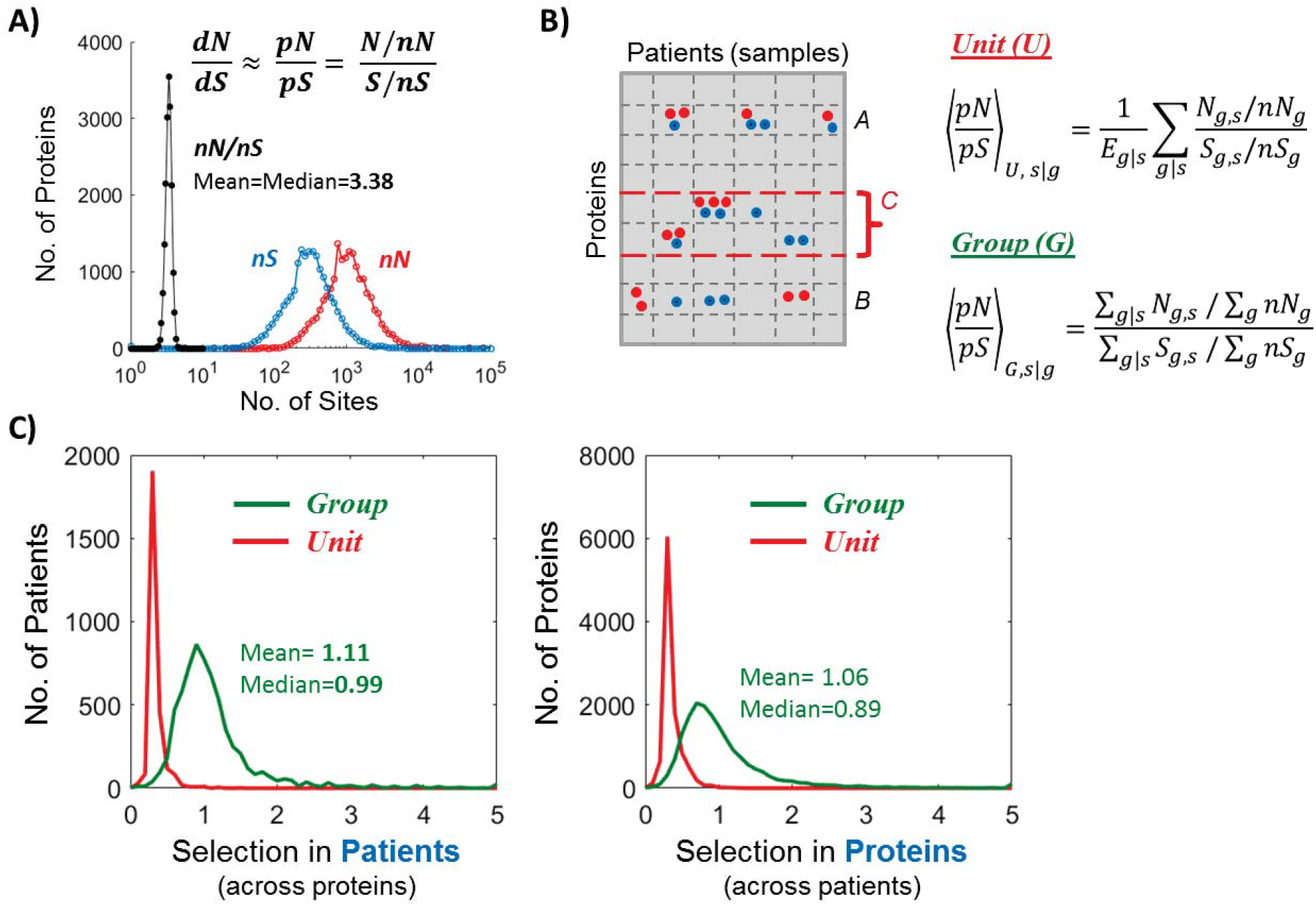
Integrative Measures of selection (*dN/dS*). **A)** Distribution of the number of non-synonymous (*nN*, red) and synonymous (*nS*, blue) sites, and their ratio (*nN/nS*, black) for all protein-coding genes (n=18179) as inferred by considering all alternative nucleotides in each position in each protein-coding sequence (**Methods**). Selection (*dN/dS*) in cancer is approximated by the ratio between the number of *N* mutations per *N* sites (*pN*=*N/nN*) and the number of *S* mutations per *S* sites (*pS*=*S/nS*). **B)** Illustration of the highly sparse mutation data, exemplified by (protein-sample pair) units that contain few mutations (e.g., protein ‘A’) and by the fact that in the vast majority of cases only one type of mutation (i.e., *N* or *S*) exists (e.g., protein ‘B’). Precisely, out of the 6721×18179 units (n=122181059), there are 963,048 cells with either *N* or *S* mutations, but only 35278 contain both *N* and *S* mutations. **C)** Unit-based and group-based estimates of selection. In principle, *dN/dS* at the proteome level can be estimated either by taking the average of *dN/dS* across units (U), or by considering a group (G) of genes (sum over *g*; for example the set ‘C’ of cancer-genes) or a group of samples (sum over *s*), and estimating *dN/dS* based on the total number of mutations in the group. Given the highly sparse mutation data, Unit-based estimators are highly biased and inadequate for analysis, whereas Group-based estimators are adequate, as exemplified by their distribution around unity. For analysis and comparisons across patients, we measured the selection in each patient, exploiting the statistical power of the overall distribution of all mutations across the proteome (summing over genes), providing a measure of selection acting on the entire proteome for each patient.

**Figure S3:**
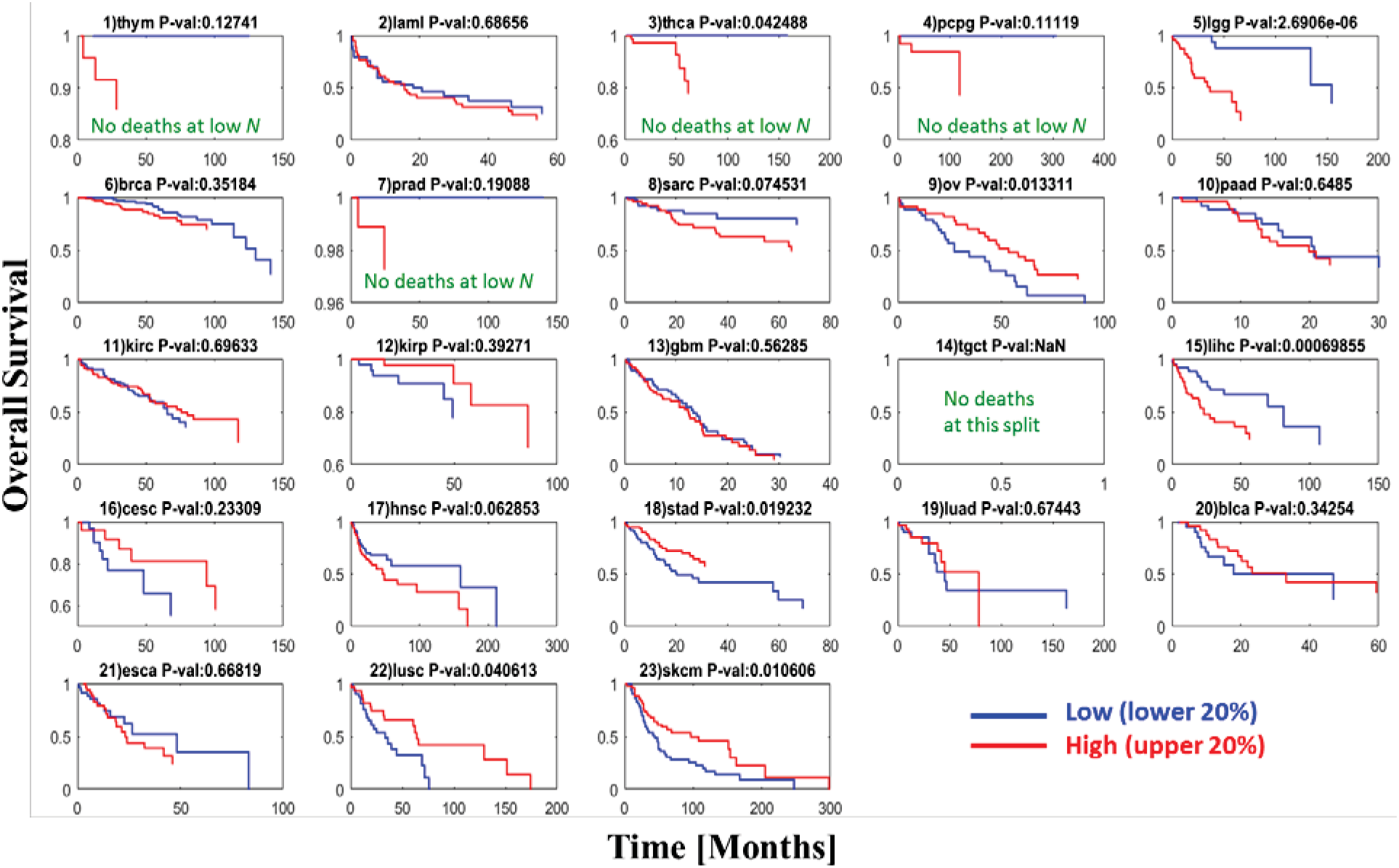
KM of mutation load (*ML*) in each cancer type. For each cancer type, survival rates were compared between patients with low (lower 20%) and high (upper 20%) number of non-silent (*N*) mutations (i.e., considering 40% of the data). Cancer types are ordered by *ML* as in **Figure 1** in the main text. At low *ML* (1-8), low number of mutations is associated with better prognosis in most cases, whereas at high *ML* (16-23), the opposite trend is observed (significant in Stad, Lusc, Skcm and to a lesser extent in Cesc and Blca). At the mutation watershed (9-15), there is no obvious trend, and Ovarian (9) and Liver (15) cancers at the edges of the watershed behave oppositely, similar to the Cox regression results summarized in **Figure 1** of the main text. Imagining the *watershed* transition as a point, it is easy to imagine that Ovarian may belong to the high *ML* cancer type and that Liver may belong to the low *ML* cancer type (main text). See **Figure 1** for the Oncotree codes of cancer types.

**Figure S4:**
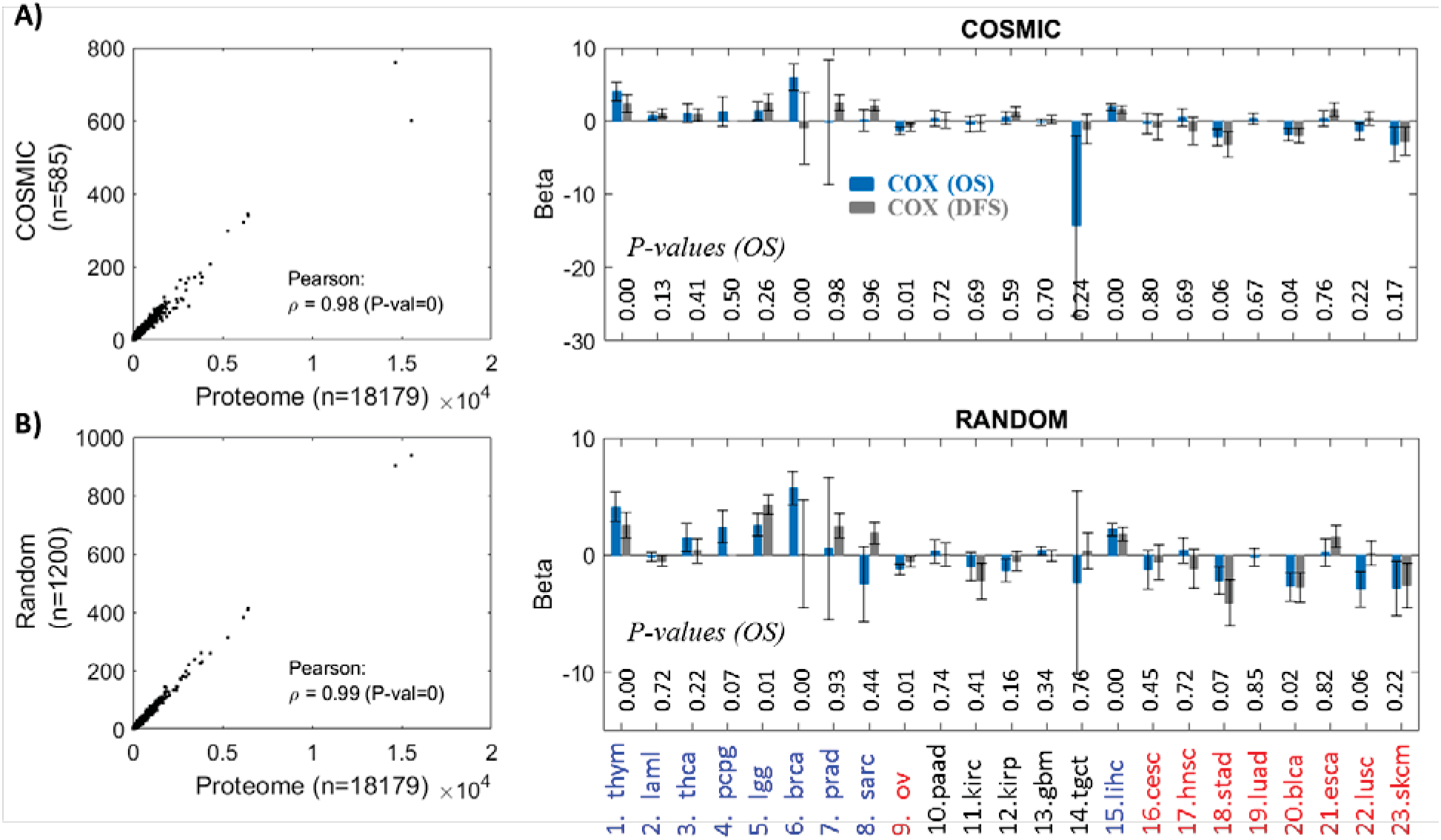
Robustness of *ML* to the gene cohort used for estimation. **A)** The correlation between *ML* estimated for the entire proteome and for cancer-genes (COSMIC) is extremely high and significant (left). Thus, it is not surprising that the pattern observed in **Figure 1** of the main text is robust to the choice of genes that are used to estimate *ML*. **B)** Similar analysis, performed for random sets of genes. A larger set of random genes had to be used to achieve a number of mutations comparable to that in the COSMIC genes which, by definition, harbor more mutations. Nonetheless, for a sufficiently large set of random genes, *ML* is highly correlated with *ML* evaluated across the entire proteome, and thus captures the transition in the clinical outcome.

**Figure S5:**
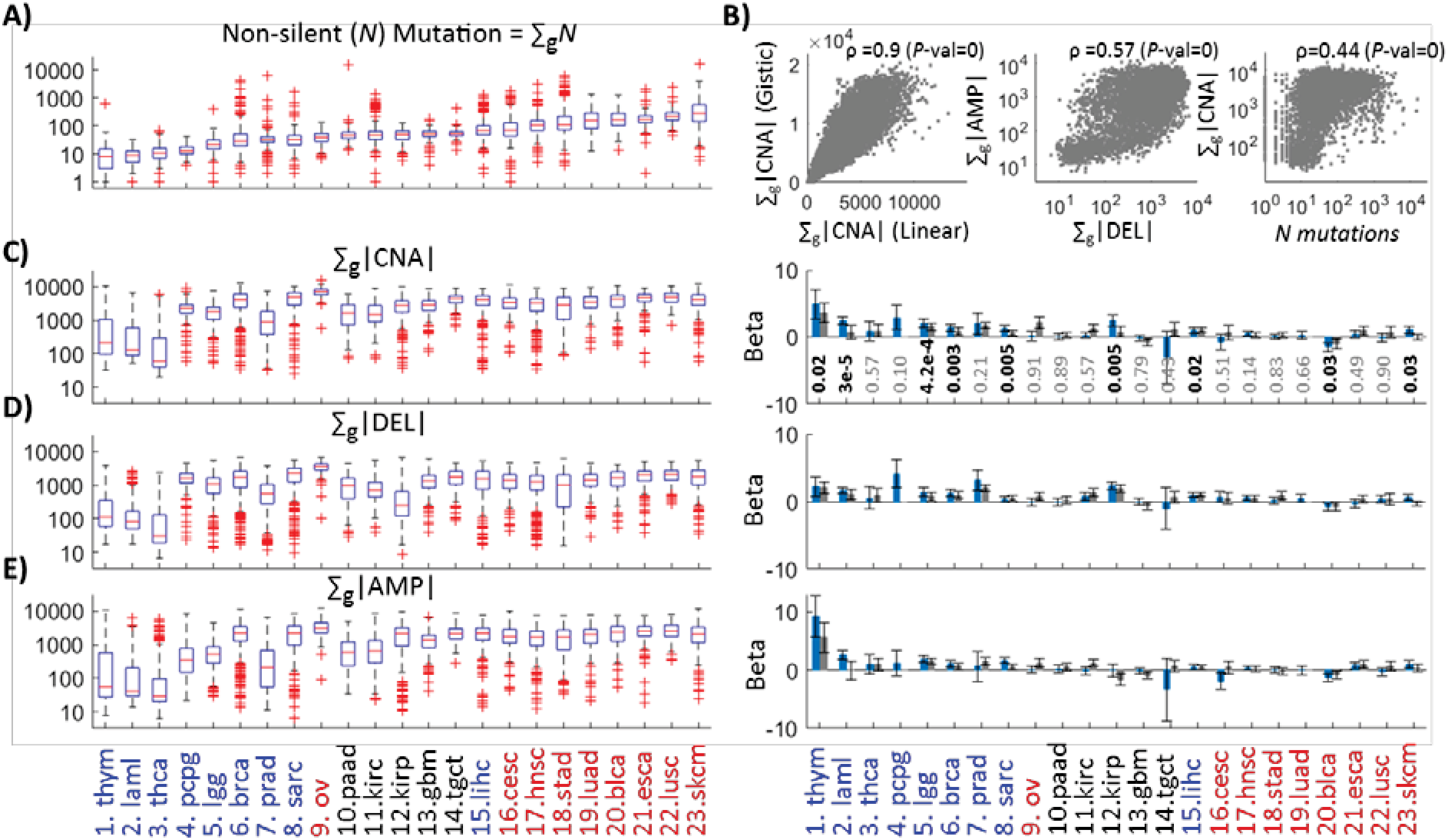
Copy number alteration (CNA) analysis and comparison to *ML*. **A)** The distribution of non-silent (*N*) mutations across cancer types as in **Figure 1** is shown, for the ease of visual comparison. **B)** Similarly to the mutation load, we estimate the overall CNA as the sum of the absolute values, using two standard measures: ‘Linear’ (a continuous variable) and ‘Gistic’ (a rounded integer variable) (**Methods**). As these measures are highly correlated they provide comparable association with survival. Continuing with the ‘Linear’ estimator, we observe that the overall CNA deletions (|DEL|) and the overall CNA amplifications (AMP) are also correlated. Also, to a lesser extent, the overall CNA (|CNA|) is correlated with the *ML*, i.e. the total number of non-silent (*N*) in each tumor proteome (Spearman correlation, p=0.44). This correlation suggests that CNA would capture a similar prognostic signal to that of *ML*. **C-E)** However, although CNA predicts pooper survival in patients harboring large CNA at low *ML*, it does not capture any transition in the clinical outcome of the type observed for *ML* in **Figure 1** of the main text. This is the case for the overall CNA (C) as well as deletions (D) and amplifications (E) each when tested separately. Complementary stratified Cox regression analysis of cancer in low and high *ML* verify these results, and show that also the copy-number DNA burden, measured as the fraction of altered genes (gain/loss) (**Methods**) behaves similarly to the overall CNA (**Table 1**).

**Figure S6:**
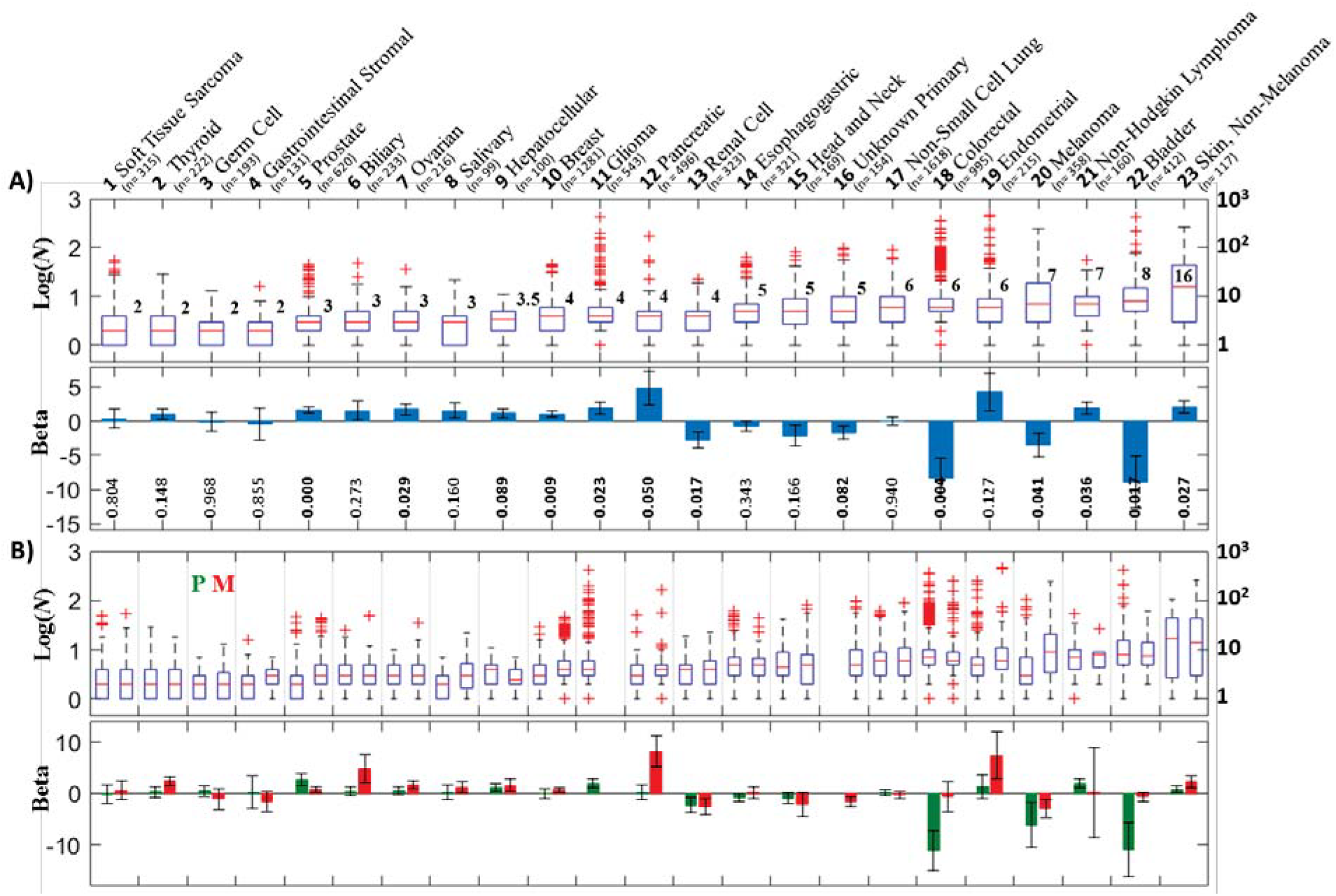
Validation by analysis of the MSK-impact-2017 cohort. A recent cohort of >10,000, which contains 43% samples from metastatic sites across >30 cancer types, was analyzed (**Methods**). Only 414 genes were sequenced in this study, for which only calls of non-silent (*N*) mutations are available. Given that our measures are apparently insensitive to the exact set of genes used to estimate *ML* (**Figure S4**), using this cohort we sought to validate our main result in **Figure 1**, that is, the existence of a transition in clinical outcome at high mutation loads (*ML*). The survival times in this cohort are provided in days intervals corresponding to the time from procedure to last follow-up. **A)** As in the main text, this analysis centered on cancer types that included at least 100 patients (one type with 99 patients was included). Discarding the sample site (i.e., including both primary and metastatic sites), we find a comparable pattern of both the mutation distribution (top) and the transition in clinical outcome, using Cox regression (bottom). **B)** The analysis was then repeated for metastatic (M, red) and primary (P, green) sites separately. Under clonal evolution, a metastatic site, by definition, should contains at least all the mutations of the corresponding primary site when taken from the same individual. The upper panel shows that, in most cases, metastatic sites indeed contain more mutations however this is not always the case because samples are taken from different individuals (top). Nonetheless, Cox regression indicates a transition in both cases although with lower significance compared to (A), presumably, because of the reduced number of patients.

**Figure S7:**
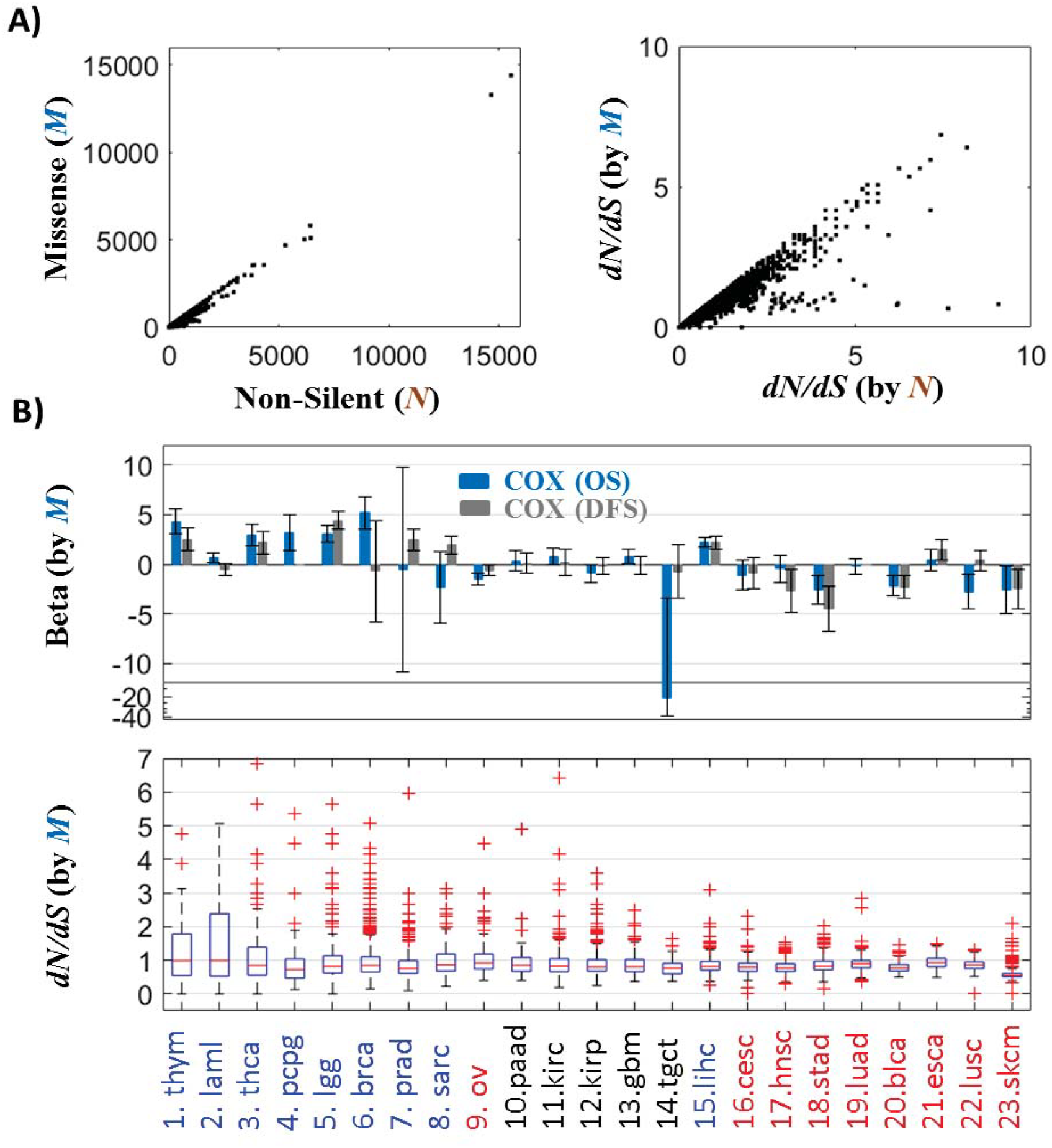
Robustness of *ML* and *dN/dS* to mutation class. The analysis presented in **Figures 1-2** of the main text was repeated for missense mutations alone. **A)** *ML* and *dN/dS* each is highly correlated for all non-silent (*N*) and missense only (*M*) mutations. **B)** The transition in clinical outcome around the mutation *watershed*, where the distribution of *ML* is flat across cancer types (**Figure 1**) is recapitulated (Top). The distributions of *dN/dS* for missense mutations only slightly shifted toward lower *dN/dS* ratios (i.e., over-estimating purifying selection), compared with the neutral evolution depicted in **Figure 1**. The heavier tails of positive selection at low *ML* and the lack thereof at high *ML*, are evident (Bottom).

**Figure S8:**
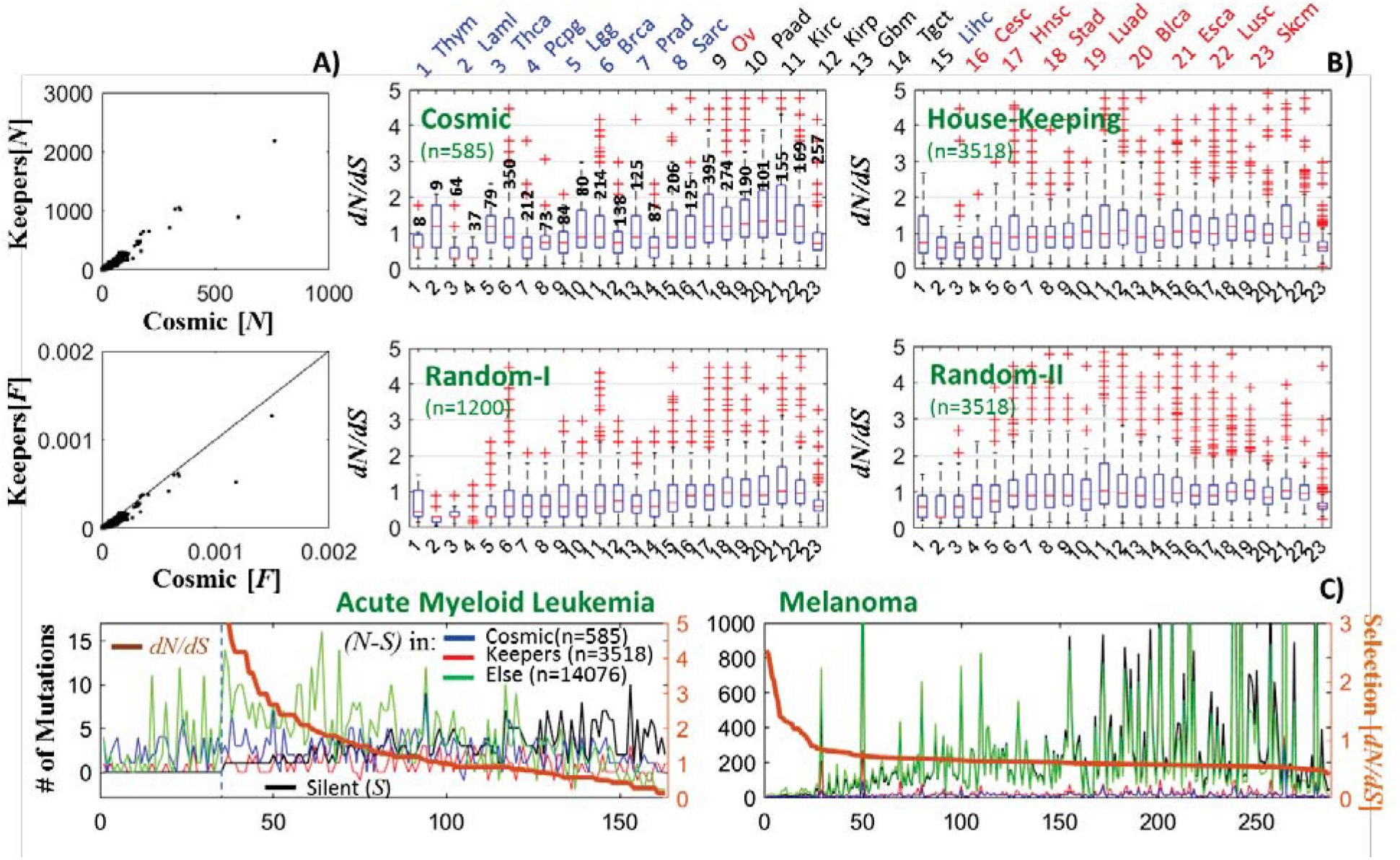
Distributions of mutations and selection (*dN/dS*) across different groups of genes. We evaluated *dN/dS* across the cohorts of patients based on the mutations in different groups of genes: Cosmic (i.e cancer-related) genes (n=585), house-keeping genes (n=3518; 145 genes that overlapped the Cosmic set were removed from the original set of 3663 house-keeping genes) and the rest of the proteome (n=14076). For each set of genes, we generated a corresponding randomized set (for the Cosmic genes, the same set as in **Figure S4** was used). **A)** Number of non-silent (*N*) mutations in Cosmic versus House-keeping genes across patients, showing the expected high correlation (**top**). Cosmic genes have higher mutation rate per unit length (*F* = *N/L* where *L* is the length of the concatenated coding sequence of the set of genes) (**bottom**). **B)** *dN/dS* distributions across patients evaluated for cancer genes and house-keeping genes (**top panels**) and respective randomized sets of genes (**bottom panels**), shown for each cancer type (ordered as in **Figure 1**). Known cancer genes displayed a much higher *dN/dS* then the respective random set, across all cancer types. In low *ML* cancers, *dN/dS* in cancer genes could be evaluated only for a small number of patients (numbers shown in figure), while in the majority of patients cancer genes harbor only *S* mutations, such that *dN/dS*=*Inf* and is discarded from analysis. Hence, **cancer genes manifest signatures of positive selection across all cancers**. In contrast, *dN/dS* ratios in house-keeping genes are distributed around unity in most cancer types, similarly to the respective randomized set. **Thus, overall, selection acts differently across different parts of the proteome, with the sum of effects leading to neutrality in most cases except at extreme *ML* (Figure 2).** In Melanoma, the cancer with the highest *ML*, purifying selection prevails in the entire proteome, and acts on each of the examined group of genes (except for cancer genes). **C)** For convenience, the distribution of mutations shown in **Figure 3** for AML and Melanoma are displayed again.

**Figure S9:**
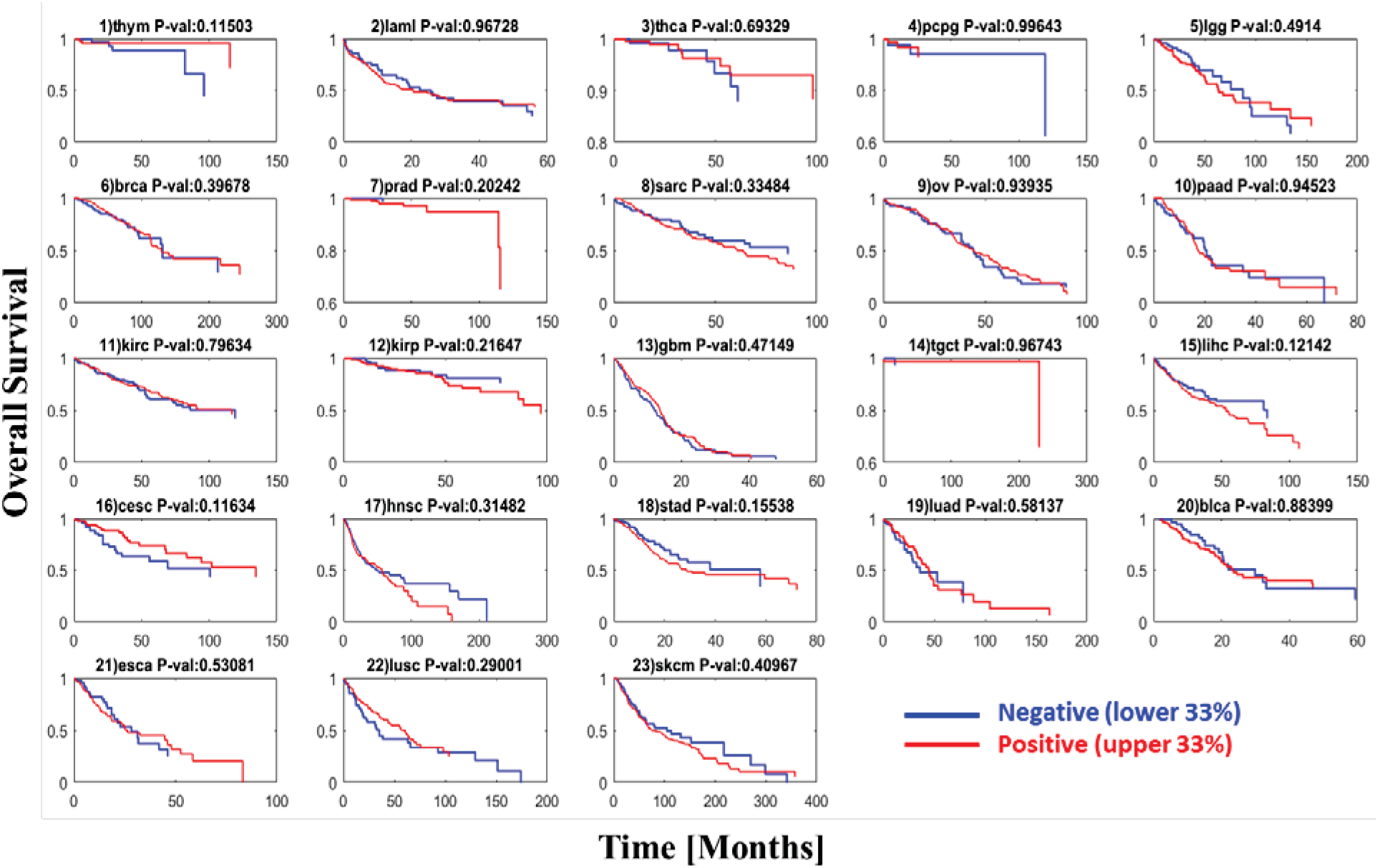
KM analysis of positive versus purifying selection in each cancer type. For each cancer type, overall survival was compared between patients with high *dN/dS* (upper 33%) indicating positive selection and patients with low *dN/dS* (lower 33%) corresponding to purifying selection. Cancer types are ordered by the mutation load (*ML*) as in **Figure 1** of the main text. No significant differences are observed, consistent with the Cox regression results shown in **Table 1** of the main text.

**Figure S10:**
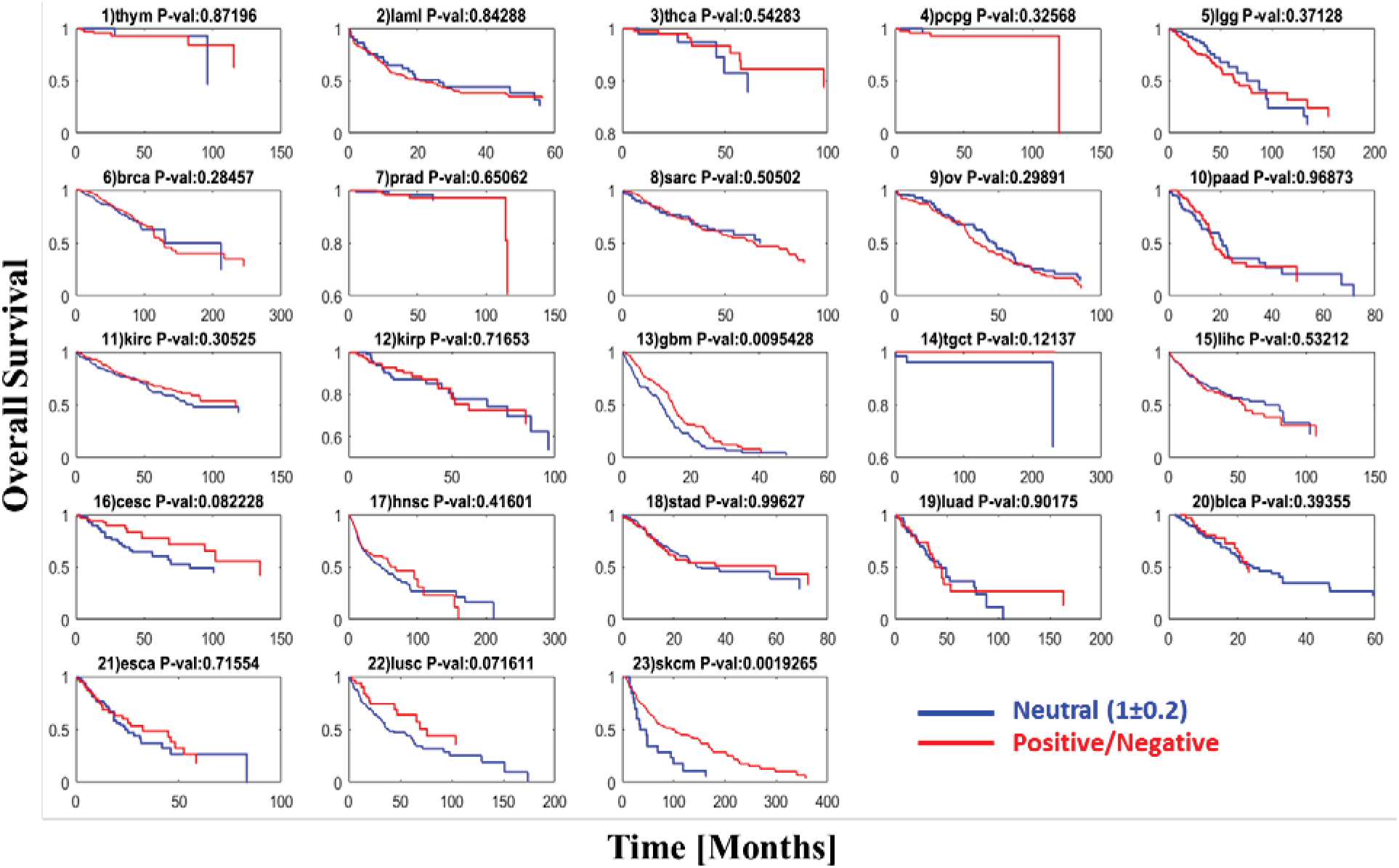
KM analysis of neutral evolution versus positive/purifying selection for each cancer type. KM for each cancer type when overall survival was compared between neutral evolution (0.8<*dN/dS*<1.2) and positive or purifying selection (*dN/dS*>1.2 and *dN/dS* < 0.8, respectively).(OS). Cancer types are ordered by the mutation load (*ML*) as in **Figures 1** of the main text. Significant differences are observed only in few a cases where neutral evolution leads to poorer prognosis (Gbm, Cesc, Lusc, Skcm). In Skcm, few patients follow neutral evolution (**Figure 2** of main text), yet their survival is much pooper than that for the rest of patients with this cancer type.

**Table S1:** Univariate versus multivariate stratified Cox regression analysis of the *ML* and confounding factors. For each tested variable the estimated scaling coefficient *β* (i.e., *HR* = *e^β^*), its standard error (SE) and the corresponding *P*-value of the stratified (by cancer type) Cox regression model are shown for overall survival (OS) and disease free survival (DFS). Univariate (U) results of **Table 1** of main text are bolded and colored (*β*>0 blue; *β*<0 red) and are compared to the results of a multivariate (M) Cox model, considering Age, Stage, Grade and overall CNA as confounding factors. Analysis is done for each of the 4 sets of cancers, corresponding to low (L) and (H) mutational load (*ML*) cancer types. Set#1 (L1, H1) corresponds to low *ML* (L1) and high *ML* (H1) cancer types, including the tips of the mutation *watershed* (i.e., 9, Ovarian; and 15, Liver), and set#2 (L2, H2) excludes these tips. In each test, variables are normalized to 0-1 within each group of cancers (**Methods**). Results of chi-square statistics, inferred by the difference in the log likelihoods of the Univariate versus Multivariate models, are shown below each test. In set#2, cancer grade is not available for the low *ML* cancers type (**Figure S1**); hence, it is removed from analysis as a confounding factor. The results verify *ML* to be the only variable which captures the transition in clinical outcome.

| Model_Variables | Cancer Set | OS |  | DFS |  |
| --- | --- | --- | --- | --- | --- |
|  |  | Beta (SE) | P-value | Beta (SE) | P-value |
| **U_ML** | **L1** | **4.41** (1.02) | **1.62e-05** | **3.12** (0.76) | **3.8e-5** |
| *M ML* | **L1** | **9.69** (2.67) | **0.00028** | **6.61** (2.85) | **0.02** |
| *M Age* | **L1** | 1.12 (0.70) | 0.11 | -0.4 (0.62) | 0.52 |
| *M Stage II* | **L1** | 0.32 (0.26) | 0.22 | **0.71** (0.21) | **0.00076** |
| *M Stage III* | **L1** | **1.05** (0.23) | **3.95e-06** | **1.01** (0.20) | **6.2e-07** |
| *M Stage IV* | **L1** | **1.52** (0.62) | **0.01364** | **1.93** (0.62) | **0.0017** |
| *M Grade II* | **L1** | 0.16 (0.32) | 0.61 | 0.35 (0.28) | 0.21 |
| *M Grade III* | **L1** | 0.22 (0.33) | 0.50 | 0.35 (0.28) | 0.21 |
| *M Grade IV* | **L1** | 0.92 (0.55) | 0.09 | 0.50 (0.53) | 0.34 |
| *M CNV* | **L1** | 0.61 (0.71) | 0.39 | 0.78 (0.63) | 0.21 |
| Chi-square (8 dof) **P-value:** |  | **0** |  | **0** |  |
| **U_ML** | **H1** | **-4.93** (1.75) | **0.0048** | **-4.24** (1.73) | **0.0145** |
| *M ML* | **H1** | **-9.12** (3.33) | **0.00624** | **-8.53** (3.9) | **0.0286** |
| *M Age* | **H1** | **2.07** (0.39) | **9e-8** | 0.76 (0.43) | 0.0753 |
| *M Stage II* | **H1** | 0.08 (0.20) | 0.71 | 0.05 (0.21) | 0.81 |
| *M Stage III* | **H1** | **0.64** (0.19) | **0.00074** | 0.35 (0.21) | 0.09 |
| *M Stage IV* | **H1** | **0.99** (0.20) | **1.4e-06** | **0.76** (0.22) | **0.000662** |
| *M Grade II* | **H1** | 0.35 (0.22) | 0.11 | 0.41 (0.25) | 0.095 |
| *M Grade III* | **H1** | 0.40 (0.22) | 0.07 | **0.54** (0.25) | **0.033** |
| *M Grade IV* | **H1** | 0.17 (1.03) | 0.87 | 0.88 (1.04) | 0.4 |
| *M CNV* | **H1** | 0.04 (0.38) | 0.92 | 0.71 (0.43) | 0.096 |
| Chi-square (8 dof) **P-value:** |  | **0** |  | **0** |  |
| **U_ML** | **L2** | **3.48** (1.46) | **0.017** | **2.81** (0.89) | **0.0015** |
| *M ML* | **L2** | **7.42** (2.34) | **0.002** | -8.73 (13.34) | 0.51 |
| *M Age* | **L2** | **3.73** (0.6) | **5.35e-10** | -0.03 (0.65) | 0.96 |
| *M Stage II* | **L2** | **0.47** (0.28) | 0.09 | 0.14 (0.27) | 0.60 |
| *M Stage III* | **L2** | **1.30** (0.29) | **5.6e-06** | **1.07** (0.26) | **3.28e-05** |
| *M Stage IV* | **L2** | **2.18** (0.35) | **6.3e-10** | **1.83** (0.36) | **4.28e-07** |
| *M CNV* | **L2** | **1.32** (0.51) | **0.001** | **0.97** (0.57) | **0.086** |
| Chi-square (6 dof) **P-value:** |  | **0** |  | **0** |  |
| **U_ML** | **H2** | **-4.79** (1.73) | **0.0057** | **-4.18** (1.73) | **0.0155** |
| *M ML* | **H2** | **-7.72** (2.05) | **0.000167** | **-5.98** (1.93) | **0.002** |
| *M Age* | **H2** | **1.96** (0.3) | **5.6e-11** | **1.37** (0.32) | **2e-05** |
| *M Stage II* | **H2** | **0.25** (0.12) | **0.0378** | 0.13 (0.14) | 0.34 |
| *M Stage III* | **H2** | **0.66** (0.11) | **1.1e-08** | **0.47** (0.13) | **0.00027** |
| *M Stage IV* | **H2** | **1.07** (0.14) | **1.5e-14** | **0.99** (0.17) | **3e-09** |
| *M CNV* | **H2** | 0.13 (0.23) | 0.57 | 0.09 (0.26) | 0.72 |
| Chi-square (6 dof) **P-value:** |  | **0** |  | **0** |  |

### Datasets

Supplemental data, provided as Microsoft Excel file, contains the samples, genes, number of mutations and clinical data, allowing fully reproducibility of the results this study.

## References

1. Knudson AG Jr. (1971). Mutation and cancer: statistical study of retinoblastoma. Proc Natl Acad Sci U S A, 68(4):820–3.

2. Cairns J (1975). Mutation selection and the natural history of cancer. Nature, 255(5505):197–200.

3. Nowell PC (1976). The clonal evolution of tumor cell populations. Science, 194(4260):23–8.

4. Sniegowski PD, Gerrish PJ, Lenski RE (1997). Evolution of high mutation rates in experimental populations of E. coli. Nature, 387:703–705

5. Feil E and Enright MC (2004). Analyses of clonality and the evolution of bacterial pathogens. Current Opinion in Microbiology, 7:308–313

6. Lang GI, Rice DP, Hickman MJ, Sodergren E, Weinstock GM, Botstein D, Desai MM (2013). Pervasive genetic hitchhiking and clonal interference in forty evolving yeast populations. Nature, 500(7464):571–4. doi: 10.1038/nature12344.

7. Gerlinger M, Rowan AJ, Horswell S, Math M, Larkin J, et al. (2012). Intratumor heterogeneity and branched evolution revealed by multiregion sequencing. N Engl J Med, 366(10):883–892. doi: 10.1056/NEJMoa1113205.

8. Ding L, Ley TJ, Larson DE, Miller CA, Koboldt DC, et al. (2012). Clonal evolution in relapsed acute myeloid leukaemia revealed by whole-genome sequencing. Nature, 481(7382):506–10. doi: 10.1038/nature10738.

9. Beerenwinkel N, Schwarz RF, Gerstung M, Markowetz F (2015). Cancer evolution: mathematical models and computational inference. Syst Biol, 64(1):e1–25. doi: 10.1093/sysbio/syu081.

10. Nik-Zainal S, Alexandrov LB, Wedge DC, Van Loo P, Greenman CD, et al. (2012). Mutational processes molding the genomes of 21 breast cancers. Cell, 149(5):979–93. doi: 10.1016/j.cell.2012.04.024.

11. Ye K, Wang J, Jayasinghe R, Lameijer EW, McMichael JF, et al. (2016). Systematic discovery of complex insertions and deletions in human cancers. Nat Med, 22(1):97–104. doi: 10.1038/nm.4002.

12. Hause RJ, Pritchard CC, Shendure J, Salipante SJ (2016). Classification and characterization of microsatellite instability across 18 cancer types. Nat Med, 22(11):1342–1350. doi: 10.1038/nm.4191.

13. Baca SC, Prandi D, Lawrence MS, Mosquera JM, Romanel A, et al. (2013). Punctuated evolution of prostate cancer genomes. Cell, 153(3):666–77. doi: 10.1016/j.cell.2013.03.021.

14. Araya CL, Cenik C, Reuter JA, Kiss G, Pande VS, Snyder MP, Greenleaf WJ (2016). Identification of significantly mutated regions across cancer types highlights a rich landscape of functional molecular alterations. Nat Genet, 48(2):117–25. doi: 10.1038/ng.3471.

15. Burrell RA, McGranahan N, Bartek J, Swanton C (2013). The causes and consequences of genetic heterogeneity in cancer evolution. Nature, 501(7467):338–45. doi: 10.1038/nature12625.

16. Hanahan D, Weinberg RA (2011). Hallmarks of cancer: the next generation. Cell, 144(5):646–74. doi: 10.1016/j.cell.2011.02.013.

17. Carter SL, Eklund AC, Kohane IS, Harris LN, Szallasi Z (2006). A signature of chromosomal instability inferred from gene expression profiles predicts clinical outcome in multiple human cancers. Nat Genet, 38(9):1043–8.

18. Birkbak NJ, Eklund AC, Li Q, McClelland SE, Endesfelder D, Tan P, Tan IB, Richardson AL, Szallasi Z, Swanton C (2011). Paradoxical relationship between chromosomal instability and survival outcome in cancer. Cancer Res, 71(10):3447–52. doi: 10.1158/0008-5472.CAN-10-3667.

19. Yuan Y, Van Allen EM, Omberg L, Wagle N, Amin-Mansour A, et al. (2014). Assessing the clinical utility of cancer genomic and proteomic data across tumor types. Nat Biotechnol, 32(7):644–52. doi: 10.1038/nbt.2940.

20. Andor N, Graham TA, Jansen M, Xia LC, Aktipis CA, Petritsch C, Ji HP, Maley CC (2016). Pan-cancer analysis of the extent and consequences of intratumor heterogeneity. Nat Med, 22(1):105–13. doi: 10.1038/nm.3984.

21. Lipinski KA, Barber LJ, Davies MN, Ashenden M, Sottoriva A, Gerlinger M (2016). Cancer Evolution and the Limits of Predictability in Precision Cancer Medicine. Trends Cancer, 2(1):49–63.

22. McGranahan N, Swanton C (2017). Clonal Heterogeneity and Tumor Evolution: Past, Present, and the Future. Cell, 168(4):613–628. doi: 10.1016/j.cell.2017.01.018.

23. Maley CC, Aktipis A, Graham TA, Sottoriva A, Boddy AM et al. (2017). Classifying the evolutionary and ecological features of neoplasms. Nat Rev Cancer, 17(10):605–619. doi: 10.1038/nrc.2017.69.

24. Bozic I, Antal T, Ohtsuki H, Carter H, Kim D, Chen S, Karchin R, Kinzler KW, Vogelstein B, Nowak MA (2010). Accumulation of driver and passenger mutations during tumor progression. Proc Natl Acad Sci U S A, 107(43):18545–50. doi: 10.1073/pnas.1010978107.

25. McFarland CD, Korolev KS, Kryukov GV, Sunyaev SR, Mirny LA (2013). Impact of deleterious passenger mutations on cancer progression. Proc Natl Acad Sci U S A, 110(8):2910–5. doi: 10.1073/pnas.1213968110.

26. Futreal PA, Coin L, Marshall M, Down T, Hubbard T, Wooster R, Rahman N, Stratton MR (2004). A census of human cancer genes. Nat Rev Cancer, 4(3):177–83.

27. Lawrence MS, Stojanov P, Polak P, Kryukov GV, Cibulskis K, et al (2013). Mutational heterogeneity in cancer and the search for new cancer-associated genes. Nature, 499(7457):214–218. doi: 10.1038/nature12213.

28. Martincorena I, Campbell PJ (2015). Somatic mutation in cancer and normal cells. Science, 349(6255):1483–9. doi: 10.1126/science.aab4082.

29. Muller HJ (1964). The relation of recombination to mutational advance. Mutat Res, 106:2–9.

30. Roberts SA, Gordenin DA (2014). Hypermutation in human cancer genomes: footprints and mechanisms. Nat Rev Cancer, 14(12):786–800. doi: 10.1038/nrc3816.

31. Williams MJ, Werner B, Barnes CP, Graham TA, Sottoriva A (2016). Identification of neutral tumor evolution across cancer types. Nat Genet, 48(3):238–244. doi: 10.1038/ng.3489.

32. Weghorn D, Sunyaev S (2017). Bayesian inference of negative and positive selection in human cancers. Nat Genet, 49(12):1785–1788. doi: 10.1038/ng.3987.

33. Martincorena I, Raine KM, Gerstung M, Dawson KJ, Haase K, Van Loo P, Davies H, Stratton MR, Campbell PJ (2017). Universal Patterns of Selection in Cancer and Somatic Tissues. Cell, 171(5):1029–1041.e21. doi: 10.1016/j.cell.2017.09.042.

34. Cooper CS, Eeles R, Wedge DC, Van Loo P, Gundem G, et al (2015). Analysis of the genetic phylogeny of multifocal prostate cancer identifies multiple independent clonal expansions in neoplastic and morphologically normal prostate tissue. Nat Genet, 47(4):367–372. doi: 10.1038/ng.3221.

35. Martincorena I, Roshan A, Gerstung M, Ellis P, Van Loo P, et al (2015). High burden and pervasive positive selection of somatic mutations in normal human skin. Science 22;348(6237):880–6. doi: 10.1126/science.aaa6806.

36. Kelly PN, Dakic A, Adams JM, Nutt SL, Strasser A (2007). Tumor growth need not be driven by rare cancer stem cells. Science, 317(5836):337.

37. Meacham CE, Morrison SJ (2013). Tumour heterogeneity and cancer cell plasticity. Nature, 501(7467):328–37. doi: 10.1038/nature12624.

38. Lynch M & Conery JS (2003). The Origins of Genome Complexity. Science, 302:1401:1404.

39. Kimura M (1962). On the probability of fixation of mutant genes in a population. Genetics, 47:713–719.

40. Lynch M (2007). The Origins of Genome Architecture. Sinauer Associates Inc (1723).

41. Koonin EV (2012). The Logic of Chance: The Nature and Origin of Biological Evolution. FT Press Science, 1st Edition.

42. Blokzijl F, de Ligt J, Jager M, Sasselli V, Roerink S, et al. (2016). Tissue-specific mutation accumulation in human adult stem cells during life. Nature, 538(7624):260–264. doi: 10.1038/nature19768.

43. Zehir A, Benayed R, Shah RH, Syed A, Middha S, et al (2017). Mutational landscape of metastatic cancer revealed from prospective clinical sequencing of 10,000 patients. Nat Med, 23(6):703–713. doi: 10.1038/nm.4333.

44. Koonin EV, Wolf YI (2010). Constraints and plasticity in genome and molecular-phenome evolution. Nat Rev Genet, 11(7):487–98. doi: 10.1038/nrg2810.

45. Eisenberg E, Levanon EY (2013). Human housekeeping genes, revisited. Trends Genet, 29(10):569–74. doi: 10.1016/j.tig.2013.05.010.

46. Campbell BB, Light N, Fabrizio D, Zatzman M, Fuligni F, et al (2017). Comprehensive Analysis of Hypermutation in Human Cancer. Cell, 171(5):1042–1056.e10. doi: 10.1016/j.cell.2017.09.048.

47. Burrell RA, Swanton C (2016). Re-evaluating clonal dominance in cancer evolution. Trends in Cancer, 2(5): 263:267. http://dx.doi.org/10.1016/j.trecan.2016.04.002

48. Tomasetti C, Vogelstein B (2015). Variation in cancer risk among tissues can be explained by the number of stem cell divisions. Science, 347(6217):78–81. doi: 10.1126/science.1260825.

49. Tomasetti C, Li L, Vogelstein B (2017). Stem cell divisions, somatic mutations, cancer etiology, and cancer prevention. Science, 355(6331):1330–1334. doi: 10.1126/science.aaf9011.

50. Polak P, Karlic R, Koren A, Thurman R, Sandstrom R, Lawrence M, Reynolds A, Rynes E, Vlahovicek K, Stamatoyannopoulos JA, Sunyaev SR. (2015). Cell-of-origin chromatin organization shapes the mutational landscape of cancer. Nature, 518(7539):360–364. doi: 10.1038/nature14221.

51. Yarchoan M, Johnson BA 3rd, Lutz ER, Laheru DA, Jaffee EM (2017). Targeting neoantigens to augment antitumour immunity. Nat Rev Cancer, 17(4):209–222. doi: 10.1038/nrc.2016.154.

52. Rosenberg SA, Aebersold P, Cornetta K, Kasid A, Morgan RA, et al (1990). Gene transfer into humans--immunotherapy of patients with advanced melanoma, using tumor-infiltrating lymphocytes modified by retroviral gene transduction. N Engl J Med, 323(9):570–8.

53. Wolchok JD1, Kluger H, Callahan MK, Postow MA, Rizvi NA (2013). Nivolumab plus ipilimumab in advanced melanoma. N Engl J Med, 369(2):122–33. doi: 10.1056/NEJMoa1302369.

54. Gao J, Aksoy BA, Dogrusoz U, Dresdner G, Gross B, Sumer SO, Sun Y, Jacobsen A, Sinha R, Larsson E, Cerami E, Sander C, Schultz N (2013). Integrative analysis of complex cancer genomics and clinical profiles using the cBioPortal. Sci Signal, 6(269):pl1. doi: 10.1126/scisignal.2004088.

55. Nei M, Gojobori T (1986). Simple methods for estimating the numbers of synonymous and nonsynonymous nucleotide substitutions. Mol. Biol. Evol. 3, 418–426.

56. Jukes TH, Cantor CR (1969). in Mammalian Protein Metabolism. Ed. Munro HN, 21–132 (Academic Press, 1969).

57. Goldman N, Yang Z (1994). A codon-based model of nucleotide substitution for protein-coding DNA sequences. Mol Biol Evol, 11, 725–736.

58. Kryazhimskiy S, Plotkin JB (2008). The population genetics of dN/dS. PLoS Genet, 4(12):e1000304. doi: 10.1371/journal.pgen.1000304.

59. Cunningham F. et al (2015). Ensembl 2015. Nucleic. Acids. Res. 43, D662–D669

60. Kaplan EL, Meier P (1958). Nonparametric estimation from incomplete observations. J Amer Statist Assn, 53(282):457–481. doi:10.2307/2281868

61. Bland JM, Altman DG (2004). The logrank test. BMJ, 328(7447):1073.

62. Cox DR (1972). Regression Models and Life-Tables. Journal of the Royal Statistical Society B, 34(2): 187–220.

